# *MetaboCensoR:* A Shiny Application for Data Filtering in Untargeted LC-MS Metabolomics to Enhance Interpretability

**DOI:** 10.64898/2026.07.02.735197

**Authors:** Ivan V. Plyushchenko, Tal Luzzatto-Knaan

**Affiliations:** Department of Marine Biology, The Charney School of Marine Sciences, University of Haifa, Israel; The Interdisciplinary Center for Metabolomics, University of Haifa, Israel

**Keywords:** LC-MS, untargeted profiling, data processing, metabolomics, R programming, software

## Abstract

Untargeted LC-MS metabolomics datasets often contain large numbers of redundant and non-informative features arising from background contaminants, multiple ion forms, poorly integrated peaks, and other low-quality signals. These features complicate downstream analysis by inflating feature space, degrading molecular networks, impeding pathway analysis, and obscuring statistically meaningful changes. Here, we present MetaboCensoR, an input-versatile Shiny application and local R package that bridges the gap between LC-MS peak picking and downstream interpretation through analyte-centric peak table filtering. The workflow integrates four fully interactive modules for blank, ion-species, quality-control, and peak-based filtering, with synchronized processing of associated spectra files. The tool was evaluated across three independent datasets covering plant extracts, human cell lines, and bacterial interactions. Across these case studies, data filtering reduced feature redundancy and improved downstream interpretation in feature-based molecular networking, pathway-level functional analysis, and differential abundance testing, while retaining known analytes. MetaboCensoR was systematically benchmarked against existing tools, demonstrating comparable ion-species annotation coverage and a favorable overall balance of target recovery, adduct precision, and feature reduction. Together, these evaluations support MetaboCensoR as a robust approach for systematic peak table curation, enhancing interpretability and analytical value of untargeted metabolomics data.

**Highlights:**

- Open-source R/Shiny GUI for untargeted LC–MS data filtering
- Enhanced interpretability across three downstream workflows
- Validated for target recovery and precision
- Benchmarked against established tools
- Robust ion redundancy removal

## 1 Introduction

Untargeted liquid chromatography–mass spectrometry (LC–MS) metabolomics processing typically begins with peak picking, resulting in feature tables generated by platforms such as xcms [1], MZmine [2], and MS-DIAL [3]. However, numerous studies report that these tables often contain up to 90% redundant and non-informative features, including background artifacts, low-quality signals, and multiple ion species (e.g., isotopes/isotopologues, adducts, multimers, neutral losses, and in-source fragments) [4–7]. Because these redundant species constitute such a large fraction of the detected signals, they severely impair downstream interpretation by inflating feature space, reducing statistical power [8,9], degrading molecular network topology [8,10], complicating both MS1 and MS2 database searches [7,11–13], and generating unknown (“dark metabolome”) features [8,14,15]. Overall, reducing feature redundancy was identified as one of the ten grand challenges in computational metabolomics in a recent global review [16].

Existing approaches address this problem only partially [9]. Background and low-quality features can be removed from peak tables by tools such as mpact [17,18] and MS-CleanR [19], while several other methods focus on grouping multiple ion features through correlation or graph-based relationships [20–23]. However, grouping approaches that lack predefined, chemically validated mass constraints are vulnerable to falsely clustering distinct, co-eluting compounds. Conversely, classical peak table annotation tools map predefined mass relationships and primarily assign ion types rather than directly removing redundant features. Examples include modules embedded within comprehensive LC–MS processing platforms such as MZmine [2] and MS-DIAL [3], and xcms-dependent packages such as CAMERA [24] and cliqueMS [25]. Tools that directly filter redundant ions remain constrained: MS-FLO [26] requires all adduct combinations to be specified individually; MS1FA [27] supports untargeted annotation of only isotopes and neutral loss relationships; khipu [28,29] focuses on isotopes and adducts; and Binner [30] combines ion annotation with broad peak grouping. Among existing approaches, MS-CleanR [19] offers a broader range of filtering options, including most major redundant ion species, but is restricted to MS-DIAL 4.0 outputs, relies on its PCE algorithm [31], and considers only features with associated MS2 spectra.

To address the fragmented landscape of feature curation, we developed MetaboCensoR: an integrated, input-versatile platform that consolidates all essential filtering steps. Designed for accessibility, it is available both as a web-based Shiny application and as a GUI-driven local R package. MetaboCensoR integrates blank filtering, comprehensive redundant ion-species reduction, QC-based filtering, and peak-based removal into a single interactive workflow with interactive threshold optimization, exportable annotation tables, and synchronized filtering of associated .mgf spectra files. Its MS-filtering strategy provides robust ion-redundancy removal through flexible criteria, conservative decision rules, and adaptive use of MS1-only or combined MS1/MS2 evidence.

MetaboCensoR performance was evaluated across three independent datasets covering plant extracts, human cell lines, and bacterial interactions. Its impact was tested in three analytical contexts: feature-based molecular networking, global pathway-level functional analysis, and univariate differential analysis. MetaboCensoR was also benchmarked against existing redundancy-reduction tools for filtering performance and ion-species annotation coverage. Across these evaluations, MetaboCensoR reduced feature redundancy while preserving genuine compounds and enhancing downstream interpretation, supporting systematic peak table curation as an important step in reliable untargeted LC-MS metabolomics analysis.

## 2 Methods

### 2.1 Computational

The MetaboCensoR application was developed within the R programming environment [32] utilizing the *Shiny* framework [33,34] to provide an interactive graphical user interface (GUI). The software employs intuitive interactive widgets, dynamic toggles, and dynamically generated tables, enabling users to conduct data curation without programming skills. The application architecture relies on robust data manipulation and visualization libraries, including *dplyr* [35], *DT* [36], and *plotly* [34]. The complete R session information and a user tutorial are available in the GitHub repository. To accommodate varying computational resources, large-scale datasets, and strict data privacy requirements, MetaboCensoR is available both as a Shiny- hosted application and as a standalone R package for local execution. For large datasets, local execution provides a more stable option by avoiding memory, timeout, and session limitations. Scalability was additionally evaluated using datasets containing up to 5,000 samples and more than 6,000 features.

Project Page: https://github.com/plyush1993/MetaboCensoR;

*Shiny* Deployment: https://plyush1993.shinyapps.io/metabocensor/;

### 2.2 Datasets

To assess the general applicability, MetaboCensoR was tested across three independent datasets. The selected datasets spanned different chromatographic modes, ionization/acquisition settings, data-processing pipelines, and replicate structures. Each dataset contained a specific set of known compounds spanning a wide dynamic range of abundances with relative peak areas from the hundreds to the millions. Using a list of target analytes defined by their chemical formulas and retention times, a broad adduct search with stringent mass and RT tolerances in MetaboAnnotation [37] was applied to search for corresponding ions, validate the accurate preservation of detected compounds after MetaboCensoR filtering, and quantify adduct-related redundancy.

A summary of the datasets is provided in Table 1, and a brief description of each dataset is given below. All code and annotated compound tables required to reproduce the three case studies are publicly available in the GitHub repository.

**Table 1.** Summary of tested datasets.

|  | Dataset ID |  |  |
| --- | --- | --- | --- |
|  | orbi | folate | inter |
| Sample Type | Plant | Human Cell Lines | Bacteria |
| Total N of samples | 9 | 37 | 28 |
| Min N of replicates | 3 | 6 | 4 |
| Chromatography | RP | HILIC | RP |
| Additives in mobile phase | HCOOH | (NH<sub>4</sub>)<sub>2</sub>CO<sub>3</sub> + NH<sub>3</sub> | HCOOH |
| MS instrument | Orbitrap | Orbitrap | TOF |
| Acquisition mode | DDA | SCAN | DDA |
| Polarity | positive | negative | positive |
| Peak Picking | MZmine | xcms | MS-DIAL |
| Total N of Known Compounds<br>(Detected by Peak Picking tool) | 153<br>(115) | 48<br>(48) | 7<br>(6) |

Case Studies, and other Supplementary code: https://github.com/plyush1993/MetaboCensoR_Examples.

#### Dataset *orbi*

The orbi dataset is an LC-MS profiling dataset described previously [38]. Methanolic extracts of the plant ashwagandha [*Withania somnifera (L.) Dunal*], together with blank samples, were analyzed. Separation was performed on a C18 column using mobile phase A composed of 100% CH_3_CN (acetonitrile) and mobile phase B composed of 100% H_2_O, both containing 0.01% HCOOH (formic acid). Data were acquired on a Thermo Q-Exactive Plus Orbitrap in positive-ion DDA mode. Raw mzML files were processed in MZmine [2] using the default UPLC-DDA preset. The spectral data and peak table, both before and after MetaboCensoR filtering, were subjected to feature-based molecular networking (FBMN) using the online workflow on the GNPS 2 platform [39], and to Ion Identity Molecular Networking (IIMN) [40]. *In silico* structure elucidation was done in SIRIUS [41] with default settings. Detailed MZmine and GNPS parameters, together with project links, are provided in the Supplementary Material, and the raw data are available in [38].

#### Dataset *folate*

The folate dataset is a human cell line metabolomics dataset, which was provided by Dr. Mariam Fokra, Dr. Nikita Sarvin, Dr. Tomer Shlomi (Technion, Israel). The Reh (CRL-8286, ATCC, USA) human cell line that was isolated from tissue from an acute lymphocytic leukemia (ALL) patient was used as a wild type (WT) control cell line. Two types of mutations were introduced to the WT in FPGS gene by CRISPR/Cas9 system. One type is the FPGS gene point mutation (PM) at the position E115K [42] identified in clinical samples diagnosed with relapsed leukemia, which is characterized by reduced FPGS activity. The other type is knockout (KO) of FPGS encoded gene. Overall, 5 cell lines were examined: WT, PM-31/37, and KO-6/22. The cell line was cultured in Roswell Park Memorial Institute medium (RPMI-1640) with a physiological level of the folic acid [43]. After washing two times with ice-cold PBS, cell pellets were resuspended in appropriate volume of MeOH–CH_3_CN–H_2_O (5:3:2 vol/vol/vol) extraction mixture. Collected samples were analyzed together with pooled QC samples. Separation was performed with a HILIC column, mobile phase A was 20 mM (NH_4_)_2_CO_3_ with 0.01% (vol) NH_3_ in H_2_O, mobile phase B was CH_3_CN. Data were collected with Thermo Q-Exactive Orbitrap in SCAN negative-mode. Raw files were converted into mzXML via MSConvert [44], and processed in xcms [1]. Target peak integration was done in El-Maven [45]. The global functional analysis [46] was performed in MetaboAnalystR (4.2.0) [47] with default settings and Mummichog p-value significance 1.0^-5^ for both raw peak table and after filtering in MetaboCensoR. Detailed xcms, El-Maven parameters and additional description are available in Supplementary Material. The converted raw data is available in the MassIVE repository under the accession number: MSV000100951.

#### Dataset *inter*

This is a bacterial interaction dataset. The study involved *Paenibacillus dendritiformis* (*Pd*), *Bacillus subtilis* NCIB 3610 (*Bs*) and the NRPS-mutated *dsrf*, *dpps*, and *dd* corresponding to surfactin-deficient, plipastatin-deficient, and double-mutant strains, respectively. A suspension of bacteria (10 μL) was inoculated from overnight cultures in LB liquid medium for microbial interactions on the nutrient plate with 0.5% bacto peptone and 1% agar. *Pd* was co-inoculated with itself, other bacteria, or 0.01 mg of commercial Surfactin at 1 cm distance to study interactions [48]. Plates were incubated at 30 °C for 24 h before analysis. Metabolites were extracted by MeOH directly from the agar cut into small pieces. Prepared extracts were analyzed together with media blank samples. Separation was performed with a C18 column, mobile phase A was 100% H2O, and mobile phase B was 100% CH3CN, both containing 0.1% HCOOH. Data were collected with a Bruker TIMS-TOF Pro 2 in DDA positive mode. Raw files were converted into mzXML via MSConvert [44], and processed in MS-DIAL [3]. The raw peak table and after filtering in MetaboCensoR were subsequently subjected to univariate analysis. Statistical power at the FDR level was estimated via pwrFDR [49]. Detailed MS-DIAL parameters and additional description are available in Supplementary Material. The converted raw data is available in the MassIVE repository under the accession number MSV000100949.

## 3 Results

### 3.1 Overview of MetaboCensoR

An overview of MetaboCensoR is presented in Fig. 1. The workflow is organized into six functional modules that collectively support peak table curation, parameter optimization, and export of filtered results.

**Fig. 1.**
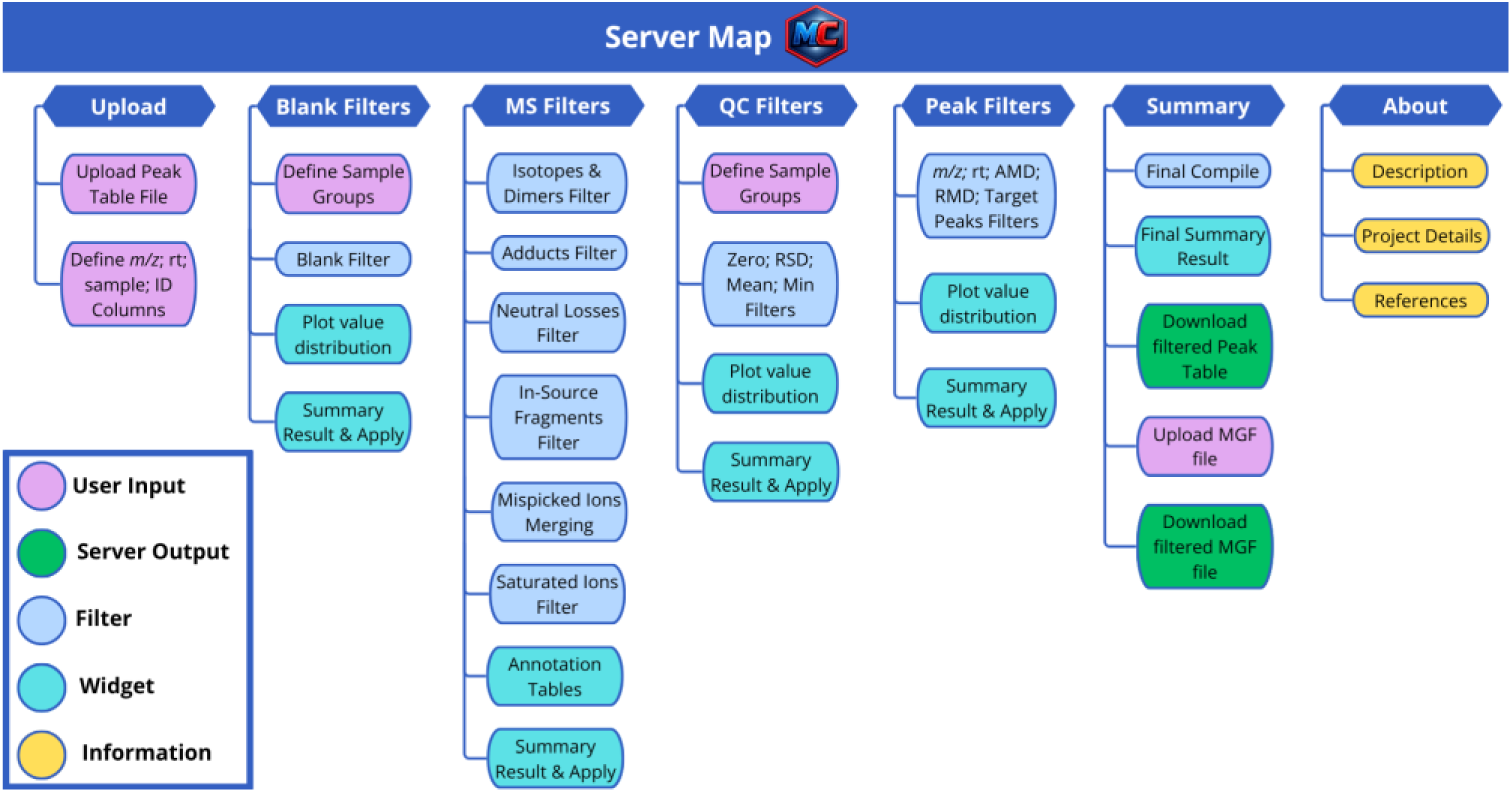
Overview of the MetaboCensoR workflow organized by application module. Colors indicate the functional role of each module.

#### Tab 1: Upload data

MetaboCensoR accepts peak tables in .csv format compatible with outputs from common peak-picking platforms, including xcms [1], MZmine [2], MS-DIAL [3]. Generic peak tables from other LC–MS processing tools can also be processed after minimal reformatting or adapted directly within the GUI. Additionally, the software accepts optional mass spectral data in .mgf format for both MS1 and MS2 levels. Users may also upload a metadata table defining sample classes, although these classes can alternatively be inferred directly from sample names.

#### Tab 2: Blank Filters

This module removes features detected in user-selected blank groups, such as media, solvent, or negative controls [9]. In **cutoff mode**, mean intensities in the selected blank groups are compared with mean intensities in the experimental groups, and a feature is retained only if the blank signal remains below a user-defined fraction of the maximum experimental signal. With the default cutoff of 0.1, the blank intensity must remain below 10% of the highest mean intensity observed in any experimental group, similar to the approach implemented in mpact [17,18] and FBMN-STATS [50]. Alternatively, in **drop any** mode, any feature detected in the selected blank groups is removed. Filtering thresholds can be adjusted interactively.

#### Tab 3: MS Filters

This module detects and collapses redundant ion species while retaining a representative feature for each analyte-related ion family [9]. It sequentially evaluates isotopes/dimer series, adducts, neutral losses (NL), in-source fragments (ISF), mispicked ions, and saturated ion artifacts. Across these steps, candidate relationships are identified using RT proximity and across-sample intensity correlation, with predefined *m/z* shifts and graph-based connectivity applied where appropriate (Fig. 2). In the illustrated example, the ^13^C isotope, [M+NH_4_]^+^, and two candidate [M+Na]^+^ ions satisfy the expected *m/z* difference and RT window relative to a representative [M+H]^+^ feature, whereas nearby ions are not assigned to the ion family. Among the two [M+Na]^+^ candidates, only one is identified as valid adduct after evaluation of across-sample intensity correlation (r > 0.8).

**Fig. 2.**
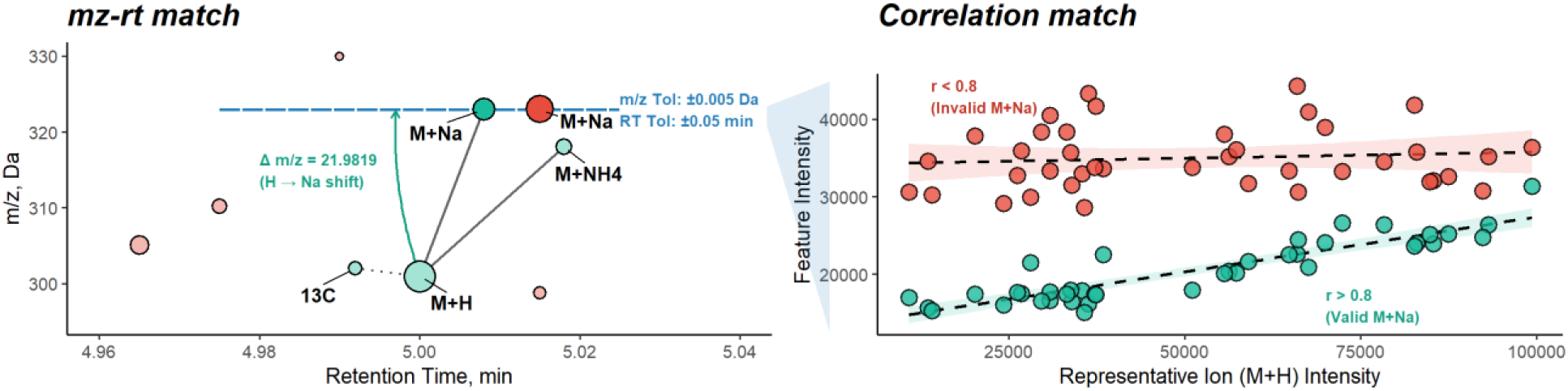
General principle of MS filtering for redundant ion-species reduction. Left: candidate relationships are identified using expected *m/z* differences and RT thresholds, linking features that satisfy the specified criteria. Right: candidate relationships are evaluated by across-sample intensity correlation; in the illustrated [M+Na]^+^ example, only the pair exceeding the correlation threshold (r > 0.80) is retained.

A brief outline of each filtering step is provided below, with full details in the Supplementary Material. Resulting annotation tables can be inspected interactively after each stage.

##### Isotopic Peak and Dimer Collapse

The algorithm searches for mass differences corresponding to ^13^C isotopes and primary dimer patterns across a user-defined range of isotope orders and charge states. Candidate pairs are filtered by *m/z* tolerance, RT proximity, and Pearson correlation, then assembled into graph-connected families, with individual feature IDs defining the graph nodes. Within each family, the feature with the highest mean intensity is retained as the representative, while the remaining nodes are removed.

##### Adduct Family Grouping

Using either a predefined [51] or user-supplied adduct library, the algorithm maps each observed *m/z* value to the theoretical neutral masses implied by the selected adduct definitions. Candidate edges are formed between features whose inferred neutral mass intervals overlap within the specified *m/z* and RT tolerances, defining the graph structure. These edges are then filtered by Pearson correlation across samples. To prevent transitive graph expansion beyond the allowed co-elution window, the remaining edges can be further restricted using strict RT control. Graph-connected components are partitioned to ensure that the maximum RT distance between any two nodes within a subnetwork does not exceed the defined RT tolerance. The resulting connected components define the final adduct families, and the most intense feature within each family is retained as the representative ion.

##### Neutral Loss Filtering

The algorithm searches for feature pairs separated by mass differences corresponding to known [27,52] or user-defined in-source neutral losses. Matching is constrained by *m/z* tolerance, RT proximity, and pairwise intensity correlation. Within each accepted pair, lower-mass fragment-like feature is removed.

##### Empirical In-Source Fragment (ISF) Collapse

In the next step, the algorithm identifies potential in-source fragments without requiring predefined mass differences. Candidate pairs are evaluated using a narrow RT co-elution window, strong correlation, and an optional intensity-ratio constraint between the putative fragment and precursor [15]. When associated MS2 spectra are available, candidate assignments can be further supported by matching the putative fragment *m/z* to fragment ions in the precursor MS2 spectrum within the specified mass tolerance and relative-intensity threshold. Within each accepted pair, the higher *m/z* feature is retained as the precursor.

##### Mispicked Ions Merging & Saturated Ions Cleaning

As a final curation step, the algorithm identifies highly correlated features within specified *m/z* and RT windows and connects qualifying pairs into graph-based components. Mispicked/misaligned features [17,18,26] are merged into the most intense representative. Saturated/ringing artifacts, associated with very high-abundance signals in TOF instruments [17,18], are removed using additional constraints based on anchor intensity, intensity ratio, and mass-difference direction. These optional filters were not used in the present study and are recommended only for poorly integrated or supersaturated signals in TOF instruments.

#### Tab 4: QC Filters

This module removes low-quality features based on quantitative reliability and signal abundance [9]. Available criteria include relative standard deviation (RSD), absolute or relative zero counts, and mean or minimum intensity, similar to the implementation in OUKS [53]. Metrics are calculated for user-selected sample groups and can be combined using logical rules such as *any*, *every*, or *pooled*. This design enables filtering based on, for example, QC reproducibility [54], missing-value patterns [55], or minimum signal abundance [56]. All thresholds are adjustable interactively.

#### Tab 5: Peak Filters

This module filters features according to peak properties such as *m/z*, retention time, and absolute or relative mass defect. These criteria can be used to incorporate prior knowledge, for example by excluding poorly-retained features, low-mass ions, or peaks outside expected mass-defect ranges [57,58]. All thresholds can be easily adjusted using dynamically updated interactive widgets. Additionally, users can retain or exclude features using a target peak list.

#### Tab 6: Summary

The final module summarizes the applied filtering steps and the resulting feature reduction. It also enables export of the filtered peak table in its original format. When an associated .mgf file is provided, the spectral file can be filtered in parallel to maintain consistency between the curated peak table and spectral data.

#### About section

An additional information section provides operating guidance, troubleshooting notes, project links, and contact details.

### 3.2 Case study 1: Improved Molecular Networking

To evaluate the efficacy of the proposed computational pipeline for profiling complex secondary metabolites, the first case study investigated natural product discovery in plant material using molecular networking, with a focus on selected chemical classes. According to the original study [38], the molecular networking results were evaluated based on their ability to successfully cluster compounds belonging to the Withanolide and Flavonol Glycoside classes into shared subnetworks. The Withanolide subnetworks, before and after data filtering, are presented in Fig. 3. Initially, we evaluated molecular-network connectivity in the raw dataset using both FBMN and IIMN. In the standard FBMN workflow, the Withanones lost connectivity with the other target compound groups, namely the Withaferins and Withanosides, and were separated into distinct subnetworks (data available in the GitHub repository). In contrast, IIMN, which incorporates additional MS1-based ion-identity relationships for potential adducts identified by MZmine, connected all three target compound groups within a single subnetwork. Based on this improvement, subsequent comparisons were performed using IIMN only. However, although IIMN restored part of the lost connectivity, its MS1-based restructuring also affected node representation. Because IIMN collapses nodes connected to the network only through MS1-based ion-identity relationships, the Withanoside V node was removed from the raw network. In addition, Withanoside III remained present but appeared only as a separated doublet (Fig. 3, left). Following MetaboCensoR filtering, network complexity was reduced while preserving structurally meaningful connectivity. The total number of connected edges decreased by 1.6-fold, from 116 to 71, while all three representative compound groups remained connected within a single subnetwork. Importantly, both Withanoside III and Withanoside V were retained and reconnected with the other Withanosides, restoring their expected structural relationships (Fig. 3, right). In addition, filtering reduced both redundancy and erroneous spectral library assignments: redundant adduct matches for Withanoside IV and Withanone were eliminated, while a node initially annotated as Withaferin A was removed as an in-source fragment and, upon further inspection, identified as originating from Sitoindoside IX.

**Fig. 3.**
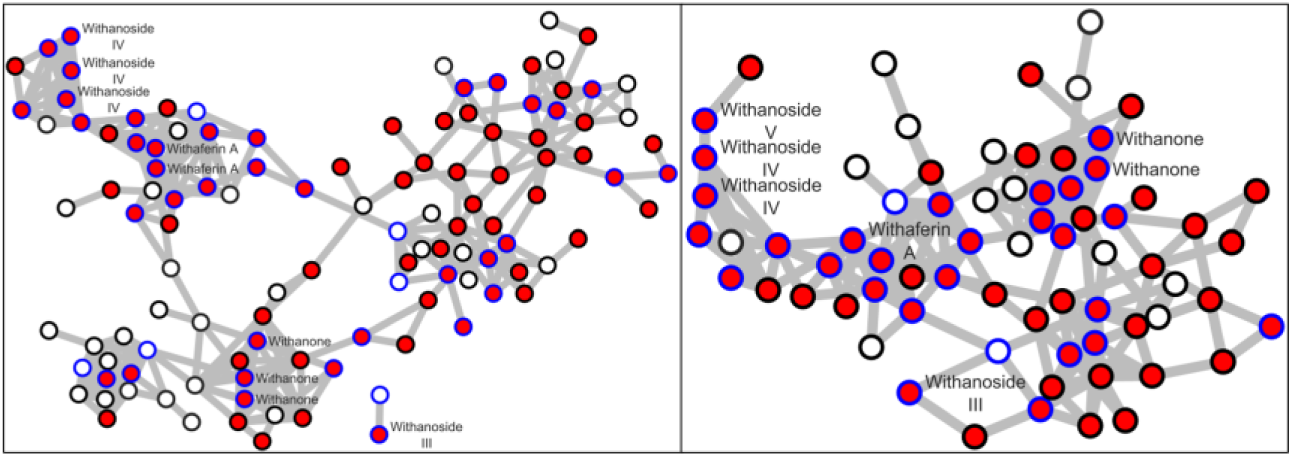
Steroids subnetwork for raw dataset (Left) and after MetaboCensoR filtering (Right). Red filling – predicted annotations for the Steroids superclass; Blue border color – matched compound with reference list of target compounds; Node labels indicate GNPS library matches.

Similarly, analysis of the Flavonoids subnetwork (Fig. 4) demonstrated a meaningful reduction in topological complexity following data curation. The total number of connected nodes decreased from 59 to 21, corresponding to a 2.8-fold reduction. Despite this reduction, all target compounds, including Rutin, Quercetin 3-O-rutinoside-7-O-glucoside, and Kaempferol-3- O-rutinoside, were preserved within the final subnetwork.

**Fig. 4.**
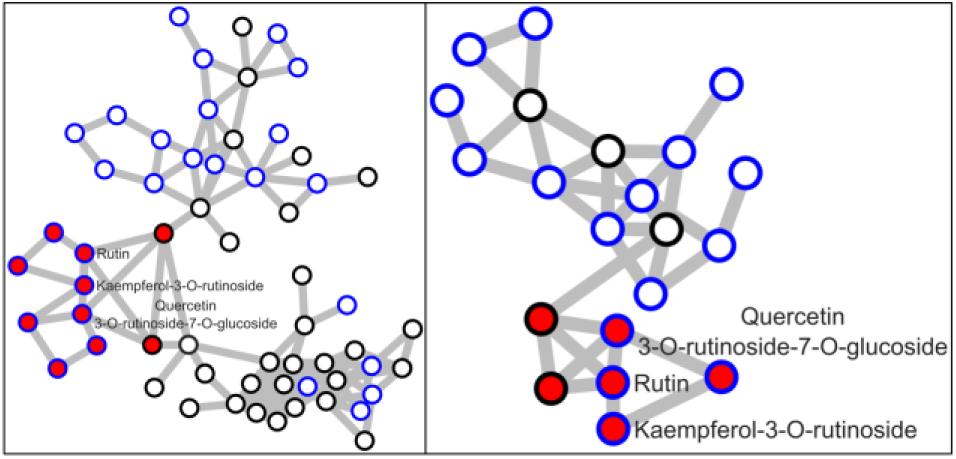
Flavonoids subnetwork for raw dataset (Left) and after MetaboCensoR filtering (Right). Red filling – predicted annotations for the Flavonoids superclass; Blue border color – matched compound with reference list of target compounds; Node labels indicate GNPS library matches.

The correctness of the data filtering was additionally validated by tracking 115 initially detected metabolites. As illustrated in Fig. 5, A, the algorithm substantially reduces multi-adduct redundancy, effectively collapsing complex clusters of up to 7 distinct features into single representative peaks, or, in total, from 315 to 147. Fig. 5, B demonstrates that the target compounds in this plant extract exhibit a vast variety of major adduct species. The retention of a relatively small subset of uncollapsed redundant adducts is primarily an artifact of the restricted experimental sample size (only 3 experimental replicates). In such small cohorts, correlations are inherently less robust, particularly for low-intensity adducts with greater relative variability, causing them to fall below the rigorous correlation threshold required for the algorithm to confidently group and collapse all corresponding features.

**Fig. 5.**
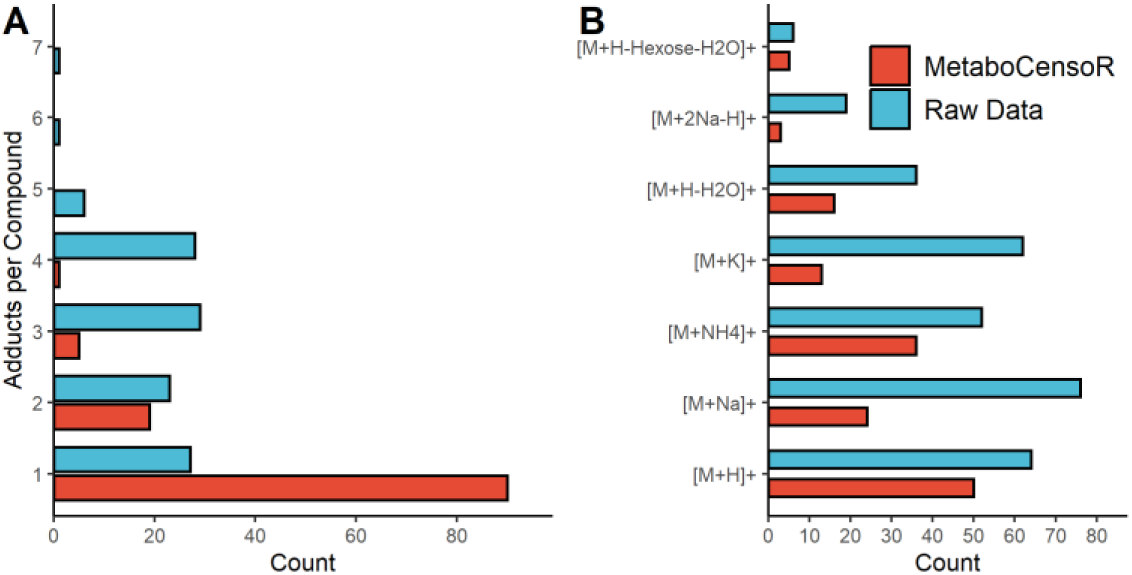
Annotation of target compounds in the raw peak table of the orbi dataset (Blue) and after MetaboCensoR filtering (Red). A – number of adducts per compound; B – adduct types.

Taken together, by systematically collapsing redundant multi-adduct clusters and removing artifactual nodes, MetaboCensoR substantially reduces network complexity without sacrificing true chemical information, streamlining downstream putative annotation and interpretation.

### 3.3 Case study 2: Improved Functional Analysis

The next case study used pathway-level global untargeted functional analysis to map metabolic alterations driven by mutations in human cell lines. Given that the induced mutations disrupt folylpolyglutamate synthase (FPGS), an essential enzyme in folate metabolism responsible for polyglutamation and subsequent intracellular retention of folic acid metabolites, significant pathway-level alterations were expected across folate-dependent processes, including nucleotide biosynthesis, SAM cycle, and Gly-Ser pathway. The results of targeted metabolomics analysis are shown in Fig. 6, where compounds are organized by their position within the pathway and compared between the mutant and WT.

**Fig. 6.**
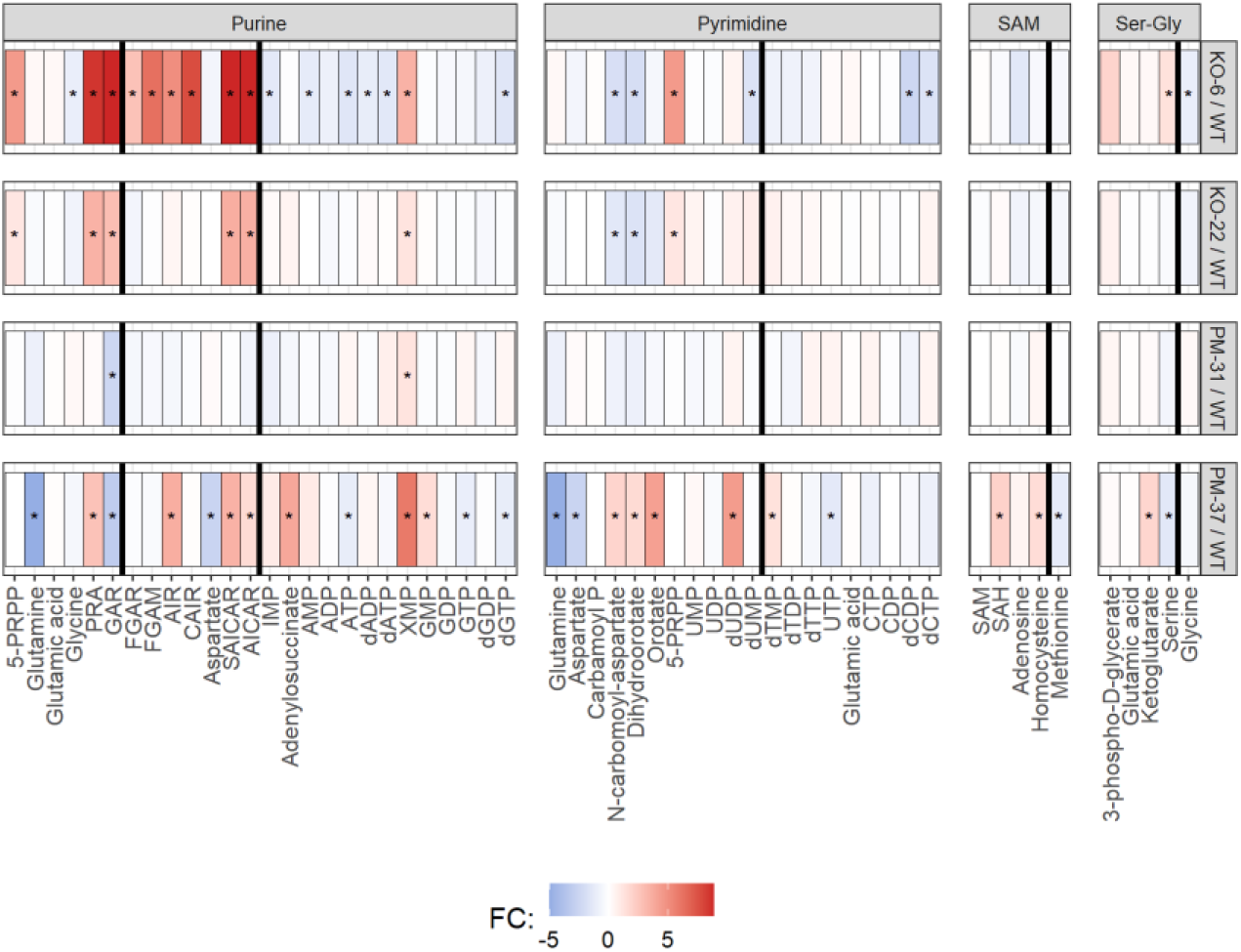
Heatmaps of the targeted metabolomics results for the studied cell lines. The colors represent the fold change (FC) value of each metabolite relative to the WT level. Star indicates statistical significance (t-test FDR < 0.05). All metabolites ordered by their appearance in pathway. Black vertical line highlights folate-dependent reactions.

Inspection of the metabolic heatmaps reveals distinct phenotypic responses across the mutant cell lines. The KO-6 line exhibits a significant inhibition in purine biosynthesis upstream of IMP, characterized by the marked accumulation of early precursors and the subsequent depletion of downstream metabolites. Conversely, the KO-22 line displays a much more moderate inhibition, with only two precursors accumulating prior to the IMP node. Within pyrimidine biosynthesis, the anticipated accumulation of uridine forms coupled with the depletion of thymidine forms was not significantly observed in these knockout lines, and other folate-dependent pathways remained largely unperturbed.

Among the point mutation lines, PM-31 predominantly mimics the WT phenotype, lacking significant metabolic shifts across the evaluated pathways. In contrast, PM-37 displays substantial, global perturbations. While the upstream inhibition of purine biosynthesis (evidenced by SAICAR and AICAR accumulation) is likely a direct consequence of the FPGS mutation, the concurrent upregulation of post-IMP purines (XMP, GMP) and diverse pyrimidine forms indicates an elevated cellular proliferation rate. This proliferative shift was also observed experimentally. Consequently, uncoupling the direct biochemical consequences of the FPGS mutation from the secondary metabolic adaptations associated with rapid proliferation in PM-37 is analytically challenging.

Ultimately, because the KO-6 line presents the most unambiguous and clearly interpretable metabolic consequence of the induced mutation, it was selected as the optimal candidate for downstream comparative analysis against the WT. The subsequent objective was to determine whether the key biological insights derived from this targeted approach could be successfully recapitulated via global untargeted functional metabolomics, utilizing the complete unannotated peak table, and to evaluate how upstream data curation impacts the accuracy of these pathway-level conclusions. The results of functional analysis for KO-6 cell line are presented in Fig. 7.

**Fig. 7.**
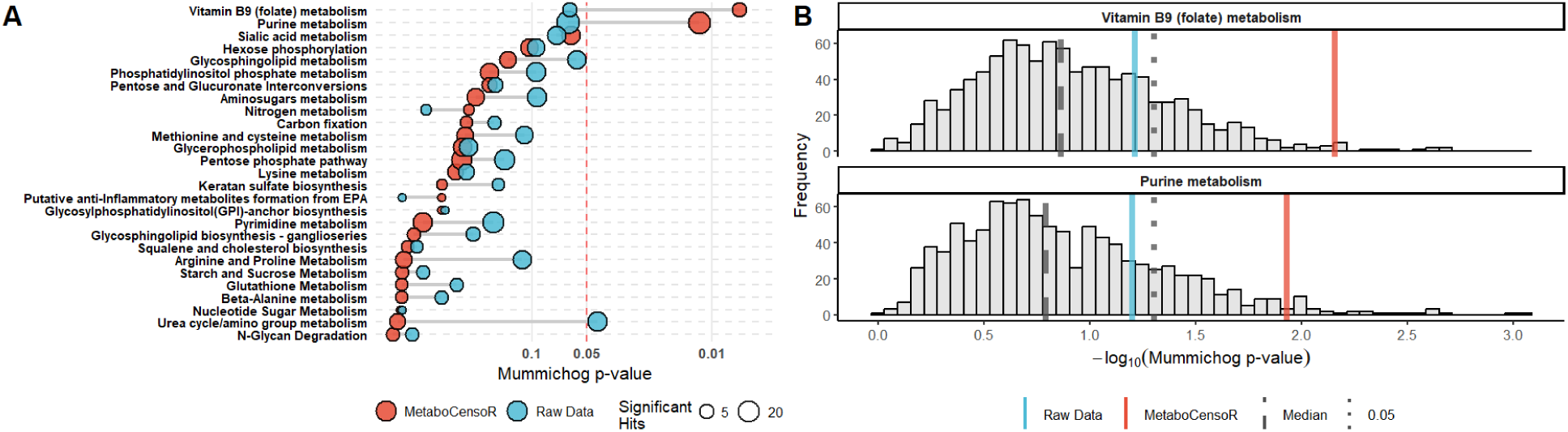
Functional analysis of KO-6 compared with WT before and after MetaboCensoR filtering. A – Mummichog p-values for enriched pathways in the raw (blue) and filtered (red) datasets; B – Distributions of Mummichog pathway significance obtained from 1,000 random size-matched subsets of the raw peak table for folate and purine metabolism. Vertical lines indicate the raw dataset (Blue), filtered dataset (Red), median of the random subsets, and p = 0.05 threshold.

As shown by the pathway enrichment analysis, MetaboCensoR filtering substantially improved the biological interpretability of the results. In the raw dataset, folate and purine metabolism appeared among the top-ranked pathways but did not reach statistical significance and competed with several unrelated pathway hits. Following filtering, these interfering hits were largely eliminated, while folate and purine metabolism became the only significantly enriched pathways, with Mummichog p-values decreasing approximately 5- and 9-fold, respectively (Fig. 7A). To determine whether this improvement could be explained simply by the reduction in feature space, the raw peak table was randomly downsampled 1,000 times to match the number of features retained after MetaboCensoR filtering, and Mummichog analysis was repeated for each subset. For both folate and purine metabolism, the filtered dataset yielded substantially lower p-values than the median of the size-matched random subsets and was located toward the extreme end of their distributions (Fig. 7B). Thus, the improved pathway enrichment cannot be explained solely by dimensionality reduction and instead reflects the selective redundant feature removal performed by MetaboCensoR.

The correctness of the data filtering was additionally validated by tracking of 48 known metabolites. MetaboCensoR successfully retained all target compounds while reducing the overall feature space by minimizing the number of adducts per compound (Fig. 8, A, in total, from 71 to 55) and substantially reducing the number of non-protonated adduct species (Fig. 8, B). A subset of uncollapsed redundant adducts remained because retention times are inherently less stable in HILIC mode, complicating the algorithm’s ability to group and collapse all corresponding features within the strict threshold.

**Fig. 8.**
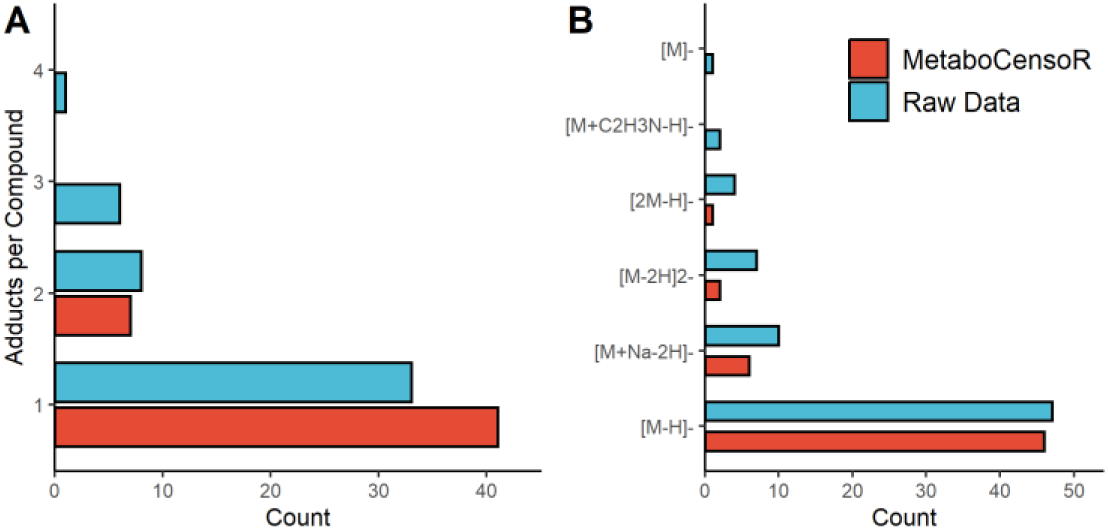
Annotation of target compounds in the raw peak table of the folate dataset (Blue) and after MetaboCensoR filtering (Red). A – number of adducts per compound; B – adduct types.

Ultimately, data curation yields a highly accurate and biologically interpretable representation of the global functional metabolic perturbations driven by the targeted genetic alterations.

### 3.4 Case study 3: Improved Statistical Hypothesis Testing

To assess the robustness of the filtering pipeline for comparative quantification, the third case study used a classical univariate statistical framework to investigate bacterial interactions and identify candidate biomarkers. According to previous findings [48], *Bs* produces Surfactin, which undergoes biotransformation by *Pd* during their interaction, generating Surfactin degradation products: a peptidic fragment (Val–Asp–Leu–Leu) and a series of lipopeptidic fragments with different carbon-chain lengths. We further introduced *Bs* mutants lacking Surfactin biosynthesis (*dsrf*), Plipastatin biosynthesis (*dpps*), or both (*dd*) to systematically examine how disruption of these pathways alters the accumulation of Surfactin degradation products during co-culture. Because Surfactin and Plipastatin are both bioactive lipopeptides, the *dpps* mutant alone was expected to show compensatory Surfactin overproduction [59] and, consequently, increased levels of Surfactin degradation products. Supplementation with pure Surfactin served as a biological positive control, while a monoculture of *Pd* was utilized as the biological negative control.

The results of the univariate statistical analysis, comparing each experimental condition against the *Pd* monoculture, are presented in Fig. 9. Data curation achieved an almost 12-fold reduction in the overall feature space, condensing the dataset from 6,027 to 509 unique features. Importantly, the target Surfactin degradation products (colored outlines) were selectively and distinctly upregulated under the expected conditions: pure Surfactin, *Bs WT*, and the *dpps* mutant. In contrast, the double mutant (*dd*) and the Surfactin-deficient mutant (*dsrf*) did not show significant overproduction of these products, consistent with the disruption of Surfactin biosynthesis in these strains.

**Fig. 9.**
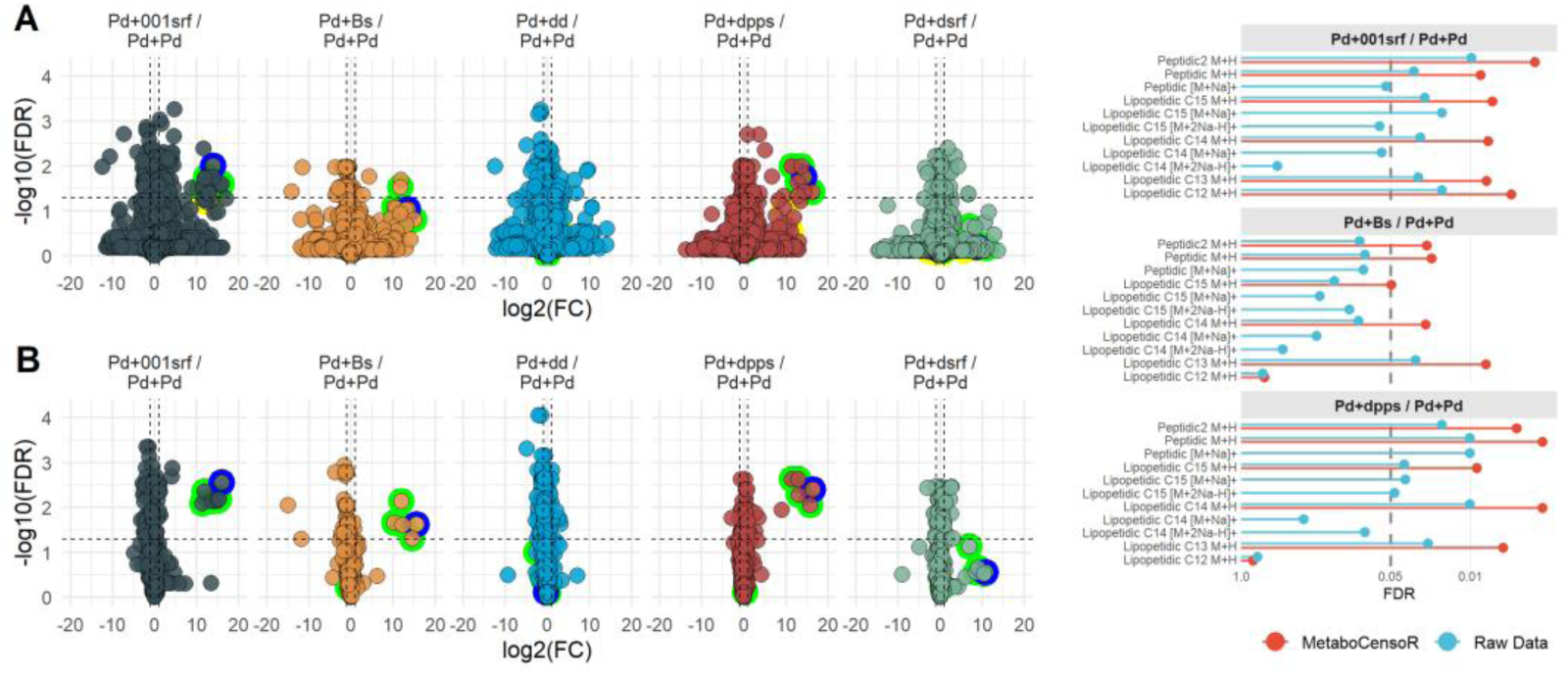
Statistical analysis of bacterial interaction dataset before and after filtering. Left: Volcano plots for all studied comparisons using the raw peak tables (top row) and the filtered peak tables (bottom row). Degradation products are highlighted by colored outlines: green – known target degradation products represented by [M+H]^+^ ions; yellow – alternative adduct forms of target compounds; blue – the previously unreported degradation product. Dashed lines indicate the significance thresholds (FDR < 0.05 and log_2_(FC) > 1 or < -1). Right: Adjusted p values (FDR) for the target compounds in the biologically relevant comparisons. Blue indicates raw data, and red indicates filtered data after MetaboCensoR processing.

Importantly, all detected degradation products, including the peptidic fragment and the C12–C15 lipopeptidic fragments, were retained after filtering and collapsed into single representative protonated ions, highlighted by green outlines in Fig. 9. In the raw data, by contrast, these same compounds were distributed across multiple alternative adduct forms, highlighted by yellow outlines. This reduction in redundancy and data dimensionality systematically improved statistical power at the FDR level. The clearest example was observed in the *Pd–Bs* / *Pd–Pd* comparison (Fig. 9, right), where filtering shifted four target degradation products below the FDR significance threshold. For these four target products, estimated absolute power increased by 26–43 percentage points.

Notably, alongside the known Surfactin degradation products, a previously unreported feature was consistently observed among the statistically significant features across the biologically relevant comparisons (blue outlines, Fig. 9). This feature was subsequently putatively annotated as a novel Surfactin-derived peptidic degradation product with *m/z* 374.2294, consistent with the Glu–Leu–Leu sequence of Surfactin (Supplementary Figs. 1–2).

Altogether, these results demonstrate that data filtering can restore the statistical power necessary to reveal biologically relevant alterations while also facilitating downstream biomarker discovery, as illustrated by the newly annotated degradation product.

## 4 Discussion

### 4.1 Assessment of MetaboCensoR Filtering

A closer examination of the filtering procedures applied to each dataset provides further insight into their overall impact. We maintained identical parameters across datasets wherever possible. This approach provides a common baseline for evaluating performance across datasets and highlights the broad applicability of MetaboCensoR under unified settings. The specific parameters for each dataset and filtering step are detailed in Table 2; notably, all shared MS filtering steps were executed using identical settings.

**Table 2.**
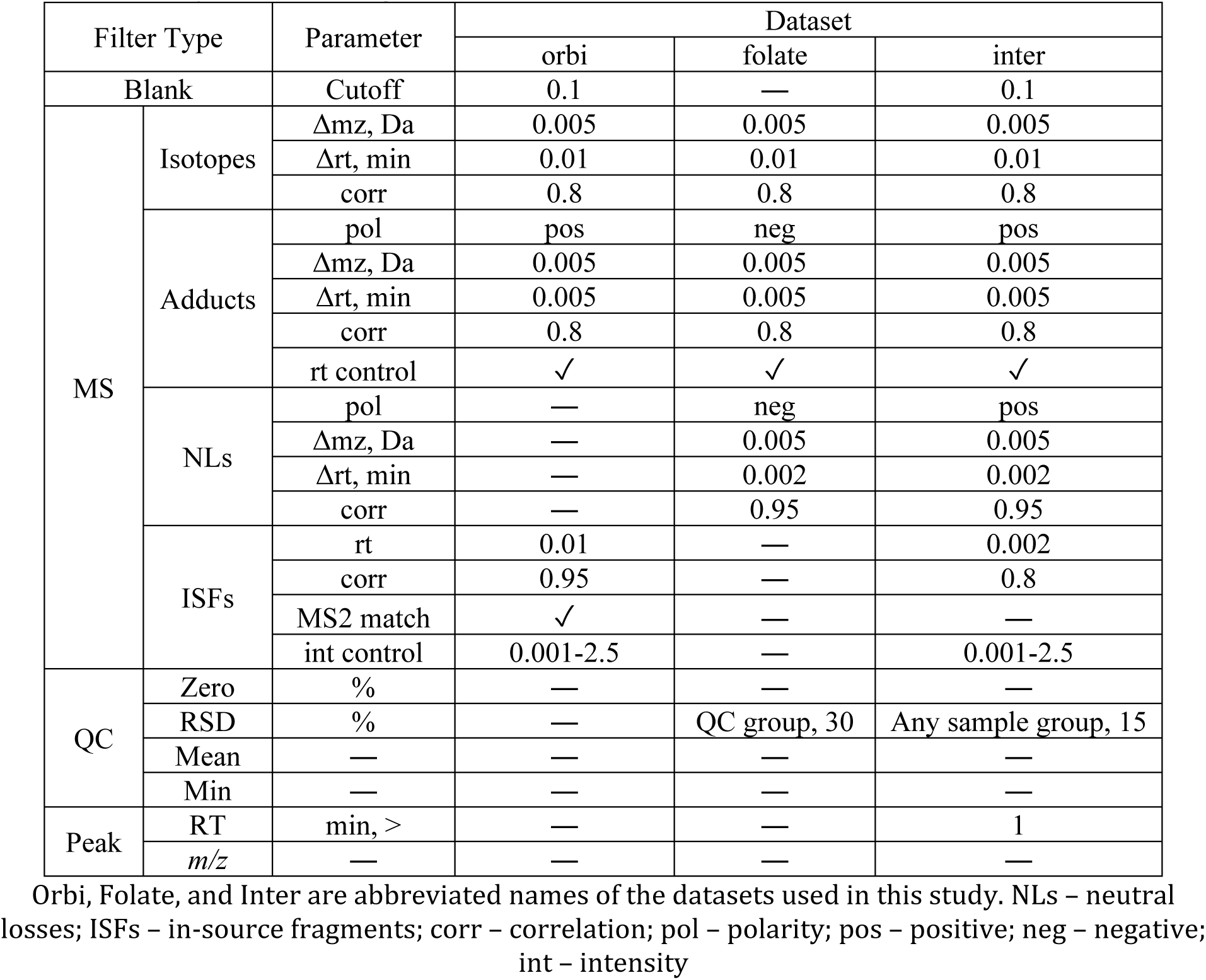
Summary of the filtering steps and applied parameters.

| Filter Type |  | Parameter | Dataset |  |  |
| --- | --- | --- | --- | --- | --- |
|  |  |  | orbi | folate | inter |
| Blank |  | Cutoff | 0.1 | — | 0.1 |
| MS | Isotopes | $\Delta m_z$, Da | 0.005 | 0.005 | 0.005 |
| | | $\Delta rt$, min | 0.01 | 0.01 | 0.01 |
|  |  | corr | 0.8 | 0.8 | 0.8 |
|  | Adducts | pol | pos | neg | pos |
| | | $\Delta m_z$, Da | 0.005 | 0.005 | 0.005 |
| | | $\Delta rt$, min | 0.005 | 0.005 | 0.005 |
|  |  | corr | 0.8 | 0.8 | 0.8 |
|  |  | rt control | ✓ | ✓ | ✓ |
|  | NLs | pol | — | neg | pos |
| | | $\Delta m_z$, Da | — | 0.005 | 0.005 |
| | | $\Delta rt$, min | — | 0.002 | 0.002 |
|  |  | corr | — | 0.95 | 0.95 |
|  | ISFs | rt | 0.01 | — | 0.002 |
|  |  | corr | 0.95 | — | 0.8 |
|  |  | MS2 match | ✓ | — | — |
|  |  | int control | 0.001-2.5 | — | 0.001-2.5 |
| QC | Zero | % | — | — | — |
|  | RSD | % | — | QC group, 30 | Any sample group, 15 |
|  | Mean | — | — | — | — |
|  | Min | — | — | — | — |
| Peak | RT | min, > | — | — | 1 |
| | $m/z$ | — | — | — | — |

Orbi, Folate, and Inter are abbreviated names of the datasets used in this study. NLs – neutral losses; ISFs – in-source fragments; corr – correlation; pol – polarity; pos – positive; neg – negative; int – intensity

The pipeline is designed to be highly adaptable to specific research needs, particularly regarding the stringency of blank and QC filtering. Through the software’s reactive interactive widgets, users can easily adjust filtering criteria in real-time. This allows researchers to verify that thresholds are not overly strict and that a reasonable number of features are retained, while still maintaining desired properties such as reproducibility, signal magnitude, and sample-to-blank ratios. For example, because MS-DIAL can occasionally generate spurious or low reproducible peak integrations [60], QC and peak filtering were specifically applied to the inter dataset to remove these artifacts.

Overall, MS filtering remains the most challenging filtering step to optimize. The isotope/dimer filtering step is relatively straightforward; given adequate LC separation, there is a low probability of detecting distinct features within a narrow RT window that differ by the ^13^C isotopic spacing, defined as: 1.00336 Da × isotope order / charge. By default, isotope detection is conservative and optimized for the regular ^13^C isotope ladder commonly observed in metabolomics peak tables. The algorithm considers isotope order 1 and charge states 1–3 and requires the first ^13^C isotope to be detected before grouping higher-order isotopes such as M+2 or M+3. However, for natural-product-rich datasets, including heteroatom-containing metabolites, an extended isotope search may be useful. For example, in the orbi dataset, increasing the isotope order to 2 with an *m/z* tolerance of 0.015 Da increased detected isotope relationships from 24 to 162. A broader tolerance should be treated as a diagnostic/refinement search and interpreted carefully, as it elevates the risk of erroneously grouping distinct co-eluting features.

Conversely, adduct filtering is more complex. Because MetaboCensoR evaluates a comprehensive default list of adducts (33 in positive mode and 16 in negative) and considers all possible combinations, distinct peaks may occasionally be grouped by neutral mass, even under strict parameters. This possibility is influenced by the accuracy and variability of *m/z* and RT measurements and by peak-integration quality, particularly for low-abundance adducts. The robustness of correlation-based grouping is additionally influenced by across-sample intensity variation and by the number and diversity of samples. For instance, the orbi dataset contained only three experimental samples, and increasing the RT tolerance from 0.005 to 0.01 min caused a feature at *m/z* 367.1897, annotated in the target list as the [M+K]^+^ adduct of C_18_H_32_O_5_, to be grouped with an unannotated ion at *m/z* 796.4136. In this case, the algorithm assigned *m/z* 367 as [M+H-H_2_O]^+^ and *m/z* 796 as [2M+3H_2_O+2H]^+^. By contrast, due to the lower RT stability of HILIC chromatography used for the folate dataset, a broader RT tolerance improved adduct grouping: increasing it to 0.1 min reduced the number of target compounds represented by multiple adducts from 7 to 1, with the remaining redundant adduct pair failing to collapse because its correlation coefficient was below 0.8. These examples illustrate the balance between conservative, reliable ion grouping and more extensive redundancy removal. The strict default parameters employed across our example datasets maximize assignment stability while minimizing data loss.

Regarding neutral loss filtering, the algorithm relies on specific mass differences (94 for positive mode and 42 for negative) without assessing combinations. Although co-eluting independent compounds may occasionally satisfy these criteria, the availability of MS2 information provides additional confidence by allowing putative neutral loss relationships to be supported by fragmentation evidence. A similar consideration applies to in-source fragment filtering, where distinguishing true fragments from independent co-eluting compounds is inherently more challenging based on MS1 information alone because of the absence of predefined mass differences. Thus, incorporation of MS2 data increases confidence in fragment assignments, whereas MS1-only filtering may enable broader feature space reduction.

Examining the quantitative impact of each filtering step (Table 3) reveals that blank filtering was responsible for the largest fraction of deleted features wherever applied. Subsequent MS filtering in the orbi and folate datasets removed a further significant portion (30–50%) of the remaining features. As expected, the number of detected multi-adducts varied extensively depending on the sample type. For instance, in the plant extracts (orbi dataset), MetaboCensoR detected thousands of redundant adducts. This is consistent with the known chemical nature of many natural product classes, which have a high propensity for forming multiple adduct species [10]. Conversely, in the folate dataset, the number of deleted isotopes and adducts was roughly equal, an outcome likely attributable to the upstream xcms processing. The inter dataset analysis exhibited the lowest number of deleted adducts, again reflecting the specific biological nature of the sample. Finally, the number of features removed during QC and peak filtering was relatively small, confirming that these steps function primarily as a final refinement stage prior to statistical analysis.

**Table 3.** Summary of the reduction in feature count following data filtering.

| Filter Type | Dataset |  |  |
| --- | --- | --- | --- |
|  | orbi | folate | inter |
| Raw | 6167 | 3201 | 6027 |
| Blank | -3570 | — | -4556 |
| MS<br>(Iso+Add+NL+ISF) | -1358<br>(24+1306+NA+28) | -1003<br>(529+438+36+NA) | -429<br>(17+184+20+208) |
| QC | — | -271 | -362 |
| Peak | — | — | -171 |
| Final | 1239 | 1927 | 509 |

Orbi, Folate, and Inter are abbreviated names of the datasets used in this study. Iso – isotopes; Add – adducts; NL – neutral losses, ISF – in-source fragments, NA – not applied

In summary, we recommend utilizing the preset universal settings and consistently applying blank filtering (when applicable), followed by isotope and adduct search. These steps are primarily responsible for eliminating feature redundancy and can be applied with relatively high confidence without extensive biological data loss, as validated across three independent datasets. For fragment filtering, when associated MS2 data are available, we recommend applying only the in-source fragment search, as it covers both neutral loss and other fragment relationships; for MS1-only datasets, we recommend applying the neutral loss search instead, as predefined mass differences provide a more constrained and robust criterion. Where supported by the study design, QC and peak filtering can be applied to further decrease dimensionality. In addition, MS-filtering thresholds, particularly for adduct grouping, can be relaxed to remove a larger number of redundant features, while retaining the most abundant ion as the representative within each accepted family; however, less stringent settings should be used cautiously because they may increase the risk of incorrect grouping.

An inherent limitation of MetaboCensoR is its dependence on the quality of upstream peak picking and on the accuracy and variability of *m/z* and RT determination. Larger-than-expected deviations or inaccurate peak integration, particularly for low-intensity features, may lead to incorrect feature assignments. Additional uncertainty may arise when the number of samples is small or when across-sample intensity patterns are insufficiently informative or highly variable, as these conditions can reduce the robustness and discriminatory power of correlation estimates.

### 4.2 Benchmarking of MetaboCensoR Performance

Beyond the primary role in redundancy reduction, MetaboCensoR can also serve as a complementary annotation source to identify putative isotopes, adducts, and fragments through dedicated, interactive data tables. To evaluate this annotation capability, MetaboCensoR was benchmarked against CAMERA [24], MS1FA [27], nontarget [51], and the built-in module of MZmine [2], across the three studied datasets (Fig. 10). For these comparisons, equivalent parameter settings were used wherever possible. MetaboCensoR demonstrated broadly comparable annotation performance across all test cases. Some reduction in coverage compared to the nontarget package is directly attributable to nontarget’s more permissive, correlation- agnostic annotation strategy. Importantly, because MetaboCensoR is designed primarily for redundancy reduction rather than definitive ion-species assignment, it does not attempt to select a single most probable adduct annotation for every feature. Instead, all plausible assignments are retained. Accordingly, the neutral-mass-based algorithm does not always uniquely distinguish between compatible ion-form assignments, such as singly charged, multiply charged, multimeric, and solvent/adduct-related forms. Thus, a single feature may receive several plausible annotations simultaneously (e.g., [M+H]^+^, [M+2H]^2+^, [2M+H]^+^, or [M+AcN+H]^+^). This conservative strategy preserves potential relationships between ion species and minimizes the risk of missing valid connections, although manual verification of MS1/MS2 spectra may still be required for exact adduct assignment. The extent of ion species connection and removal can be further optimized by adjusting threshold stringency and refining the lists of considered adducts.

**Fig. 10.**
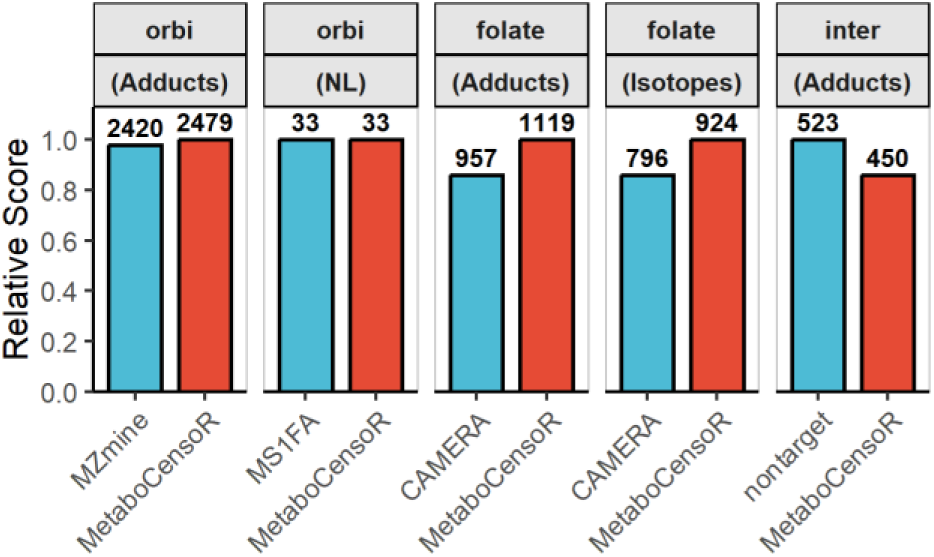
Annotation coverage of MetaboCensoR in comparison with existing tools. Numbers above the bars indicate the number of annotated features, and relative scores are normalized to the maximum value within each comparison.

To facilitate this, MetaboCensoR provides a summary of all detected adducts within a dedicated table widget, allowing iterative optimization of the adduct list based on frequency of occurrence.

As a final evaluation, the overall filtering performance of MetaboCensoR was benchmarked using the orbi dataset against existing tools designed for LC–MS redundancy reduction: khipu [28,29], mpactR [17,18], MS-CleanR [19], and Binner [30]. The evaluated tools differ substantially in their input requirements and available filtering functions (Table 4). For each benchmarked tool, all applicable filtering options listed in Table 4 were included in the comparison. Wherever possible, equivalent filtering steps and comparable parameter settings were applied to MetaboCensoR and the corresponding benchmark tool; full details are provided in the Supplementary Material. Because individual tools impose different input and processing requirements, comparison conditions were adapted accordingly, while identical input data were used within each head-to-head comparison.

**Table 4.** Functional capabilities of benchmarked tools and MetaboCensoR.

| Software | Input | Blank | MS | QC | Peak | Spectra Handling |
| --- | --- | --- | --- | --- | --- | --- |
| khipu | Custom | – | Iso, Add | – | – | – |
| mpact, mpactR | mzMine, Progenesis, MetaboScape, MS-DIAL, Custom | + | Iso, MP, SI, ISF | RSD | – | – |
| MS-CleanR | MS-DIAL 4.0 (MS2) | + | Iso, Add, NL, ISF, MP | RSD | RMD | + |
| Binner | Custom | – | Iso, Add, NL, ISF | RSD, Zeros | – | – |
| MetaboCensoR | mzMine, xcms, MS-DIAL, Custom, GUI adaptive | + | Iso, Add, NL, ISF, MP, SI | Mean, Min, RSD, Zeros | $m/z$, RT, RMD, AMD, Peak List | + |

RSD – relative standard deviation, Iso – isotopes; Add – adducts; NL – neutral losses, ISF – in-source fragments, MP – mispicked ions, SI – saturated ions, RMD – relative mass defect, AMD – absolute mass defect

Filtering performance was evaluated using three complementary metrics: Target Compound Recovery, Adduct Precision, and Relative Feature-Space Reduction (Fig. 11).

**Fig. 11.**
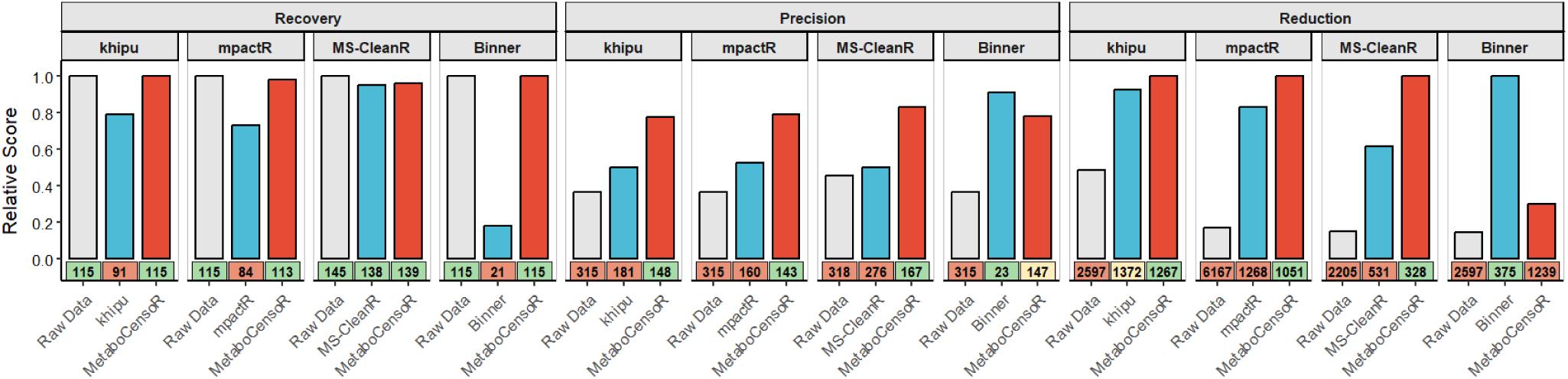
Benchmarking of MetaboCensoR against existing redundancy reduction tools using Recovery, Precision, and Reduction metrics. Relative scores were normalized to the best performing result within each comparison, with higher values indicating better performance. Numbers below the bars show absolute counts; background colors indicate performance relative to the best result: green, >0.95; yellow, 0.85-0.95; red, <0.85. Because the tools differ in input requirements and filtering capabilities, comparison conditions varied between tools, while identical input data and matched settings were used within each comparison.

Target Compound Recovery was calculated as the ratio between the number of unique known compounds retained after filtering (*N_Filt_*) and the number detected in the initial dataset (*N_Raw_*): *Recovery* = *N_Filt_*⁄*N_Raw_*. A value of 1.0 therefore indicates that all initially detected known compounds were preserved after filtering.

Adduct Precision was calculated as the ratio between the number of unique annotated known compounds (*N_Uni_*) and the total number of corresponding annotated ions/adducts (*N_Tot_*):

*Precision* = *N_Uni_*⁄*N_Tot_*. This metric reflects the degree of adduct redundancy remaining after filtering, with a value of 1.0 indicating that only one representative ion remained for each unique compound.

Relative Feature-Space Reduction was calculated as the ratio between the number of features in the most reduced peak table within each head-to-head comparison (*N_min_*) and the number of features remaining after filtering by the evaluated tool (*N_tot_*): *Reduction* = *N_min_*⁄*N_tot_*. A value of 1.0 indicates the greatest feature-space reduction observed within that comparison.

The benchmarking revealed clear differences across the three performance metrics (Fig. 11). For Target Compound Recovery, MetaboCensoR consistently retained >95% of initially detected known compounds across all comparisons, whereas greater losses were observed for several benchmarked tools, most notably Binner, which retained only 21 of 115 compounds, likely as a consequence of its aggressive feature-grouping strategy. For Adduct Precision, MetaboCensoR substantially outperformed khipu, mpactR, and MS-CleanR, indicating more effective removal of adduct redundancy. Binner achieved the highest precision, but this coincided with extensive target loss. A similar pattern was observed for Relative Feature-Space Reduction: MetaboCensoR achieved the greatest reduction in comparisons with khipu, mpactR, and MS-CleanR, whereas Binner produced a stronger reduction at the expense of compound recovery. Overall, the benchmarking highlights that effective feature filtering requires a balance between redundancy reduction and preservation of genuine analytes. Across the evaluated tools and metrics, MetaboCensoR provided a favorable balance of high target recovery, adduct precision, and feature-space reduction, supporting efficient data curation with minimal loss of relevant information.

## 5 Conclusions

MetaboCensoR provides an integrated, analyte-centric framework for systematic peak- table curation in untargeted LC–MS metabolomics. By combining blank removal, comprehensive redundant ion-species reduction, QC-based filtering, and peak-level filtering within a single interactive workflow, the platform addresses multiple sources of feature redundancy while retaining flexibility for different datasets and experimental designs. Across three independent datasets, MetaboCensoR substantially reduced feature-space complexity while preserving known compounds and improving downstream interpretation in molecular networking, pathway-level functional analysis, and univariate statistical testing.

Benchmarking against existing tools demonstrated broadly comparable ion-species annotation coverage and, in separate filtering-performance comparisons, a favorable balance between target compound recovery, adduct precision, and relative feature-space reduction, without substantial loss of genuine analytes.

From a functional perspective, MetaboCensoR combines broad input compatibility with the most comprehensive range of filtering options among the evaluated tools, supports both MS1-only and combined MS1/MS2 workflows, and provides interactive parameter optimization, ion-species annotation, and synchronized processing of associated spectral files. Its GUI-guided design enables researchers to incorporate systematic peak-table curation into existing metabolomics workflows with minimal programming effort while retaining full control over filtering criteria and thresholds. Evaluation across the three studied datasets also established a consistent set of recommended parameter settings that can serve as a practical starting point for routine analysis. Together, these capabilities provide an accessible and reproducible bridge between LC–MS peak picking and downstream metabolomics analysis. Taken together, the findings indicate that MetaboCensoR offers a robust framework for systematic peak-table curation, reducing feature redundancy while preserving genuine analytes and improving the interpretability and analytical value of untargeted LC–MS metabolomics data.

## Data and Software Availability

The MetaboCensoR source code is available at https://github.com/plyush1993/MetaboCensoR. The deployed web application is available at https://plyush1993.shinyapps.io/metabocensor. Case-study scripts, all computational code, figure-generating code, benchmarking and scalability data, session info, tutorial, input files, and processed outputs are available at https://github.com/plyush1993/MetaboCensoR_Examples. The previously published orbi dataset was obtained from the study by Houriet et al. [38]. The folate dataset (folate) is available in the MassIVE repository under accession MSV000100951 and DOI 10.25345/C51J97N5K. The bacterial interaction dataset (inter) is available in the MassIVE repository under accession MSV000100949 and DOI 10.25345/C5930P825.

## Appendix A. Supplementary Material

Detailed MS filtering algorithms; data-processing parameters and GNPS/Cytoscape projects for the orbi dataset; sample preparation, acquisition settings and data processing parameters for the folate dataset; sample preparation, acquisition settings and data processing parameters for the inter dataset; MS/MS putative annotation of the previously unreported peptidic degradation product in Supplementary Figs. 1-2; parameter settings used for benchmarking the evaluated tools (PDF)

## CRediT authorship contribution statement

The manuscript was written through contributions of all authors. All authors have given approval to the final version of the manuscript. IVP: conceptualization, methodology, software, investigation, writing; TLK: supervision, project administration, data interpretation, funding acquisition, writing.

## Declaration of generative AI and AI-assisted technologies

During the preparation of this manuscript, the authors used ChatGPT (OpenAI) and Gemini (Google) chatbots to edit text for clarity and readability, and to refine R scripts. The authors reviewed and edited all outputs and take full responsibility for the content of the published article, including all associated data and code.

## Acknowledgments

This project was funded by Haifa University Institutional Postdoctoral Scholarship for 2025-2026 academic year awarded to IVP. The authors thank Dr. Mariam Fokra, Dr. Nikita Sarvin, and Dr. Tomer Shlomi (Technion–Israel Institute of Technology, Israel) for providing the dataset used in this work.

