## Supplementary Material for "*MetaboCensoR:* A Shiny Application for Data Filtering in Untargeted LC-MS Metabolomics to Enhance Interpretability"

### Contents

Detailed description of MS Filtering Steps ... 4
- Isotopic Peak and Dimer Collapse ... 4
- Adduct Family Grouping ... 4
- Neutral Loss (NL) Filtering ... 5
- Empirical In-Source Fragment (ISF) Filtering ... 5
- Mispicked Ion Merging ... 5
- Saturated (Ringing) Ion Cleaning ... 5

Orbi dataset ... 7
- Data Processing ... 7
  - MZmine ... 7
  - GNPS ... 9
- Data Availability ... 9

Folate dataset ... 10
- Human Cell Lines ... 10
- Sample Preparation ... 10
- LC-MS Analysis ... 11
- Data Processing ... 11
  - El-Maven ... 11
  - xcms ... 11
- Statistical Analysis ... 12
- Data Availability ... 12

Inter dataset ... 13
- Bacterial Strains ... 13
- Sample Preparation ... 13
- LC-MS Analysis ... 13
- Data Processing ... 14
  - MS-DIAL ... 14
- Statistical Analysis ... 15

### Detailed description of MS Filtering Steps

#### Isotopic Peak and Dimer Collapse

The algorithm constructs target mass shifts corresponding to $^{13}C$ isotopes ($\Delta m/z$ = 1.00336×$n/z$, up to a maximum user-defined charge $z$ and isotope order $n$) and primary dimer formations (1.00336×($n\_d$ + 0.5)). By default, $n$ = 1, $n\_d$ = 3, $z$ = 3. Candidate feature pairs falling within the target $\Delta m/z$ (default ≤ 0.005 Da) and maximum RT tolerance (default ≤ 0.01 min) are validated via Pearson correlation. Edges are retained if $r$ exceeds the user-defined threshold (default $r \geq 0.80$). Validated pairs are connected to form a network graph, with individual feature IDs defining the graph nodes, grouping all related features into a single interconnected family. Connected components are extracted, and the feature exhibiting the highest mean intensity across all samples is retained as the representative. All other nodes within the component are pruned. Isotope-dimer table used in the App is available in the GitHub.

#### Adduct Family Grouping

Relying on a predefined (derived from [1], available in the GitHub) or user-supplied library of adducts, the algorithm maps the observed $m/z$ of each feature to the theoretical neutral masses $M$ (default > 50 Da) implied by the selected adduct definitions. Candidate edges are formed between any two features whose inferred $M$ intervals overlap within the defined mass tolerance (default ≤ 0.005 Da), and RT window (default ≤ 0.005 min). An initial network graph, with individual feature IDs defining the graph nodes, is generated. Then, Pearson correlation is calculated across all samples for the surviving edges. Edges that fail to meet the user-defined correlation threshold (default $r \geq 0.80$) are severed, pruning unrelated co-eluting background noise from the adduct network. Optionally, to prevent a topological artifact where a chain of adjacent nodes cumulatively exceeds the defined co-elution RT window between its ends, the strict RT split is applied. Graph-connected components are computationally fractured to ensure the maximum RT distance between any two nodes in a sub-network strictly complies with the defined RT tolerance. The final connected components are extracted, and the node exhibiting the highest mean intensity across samples within each refined cluster is designated as the representative adduct and retained, while the redundant adduct features are removed.

#### Neutral Loss (NL) Filtering

The algorithm scans for mass differences between feature pairs to match a library of known in-source neutral losses (derived from [2,3], available in the GitHub). This search is constrained using defined thresholds for retention time (default ≤ 0.002 min), pairwise intensity correlation (default $r \geq 0.95$), and mass tolerance (default ≤ 0.005 Da). The algorithm retains the ion with the highest *m/z* in each pair, pruning the lighter fragment.

#### Empirical In-Source Fragment (ISF) Filtering

At next pass to capture unannotated or complex fragmentation, the algorithm is agnostic to exact *m/z* values. Candidate pairs are formed between any features that strictly co-elute within the narrow RT tolerance (default ≤ 0.002 min), and strong Pearson correlation (default $r \geq 0.95$). Because no theoretical mass shift anchors this step, a strict intensity ratio boundary is enforced. The ratio of the putative fragment to the precursor must fall within relevant limits (default 0.001 to 2.5, according to the study [4]). When associated MS2 spectra are available, candidate assignments can be further supported by matching the putative fragment *m/z* to fragment ions in the precursor MS2 spectrum within the specified mass tolerance (default 0.01 Da) and relative intensity threshold (default > 1%). Within each accepted pair, the feature with the highest *m/z* is assigned as the true precursor and retained, while the fragment is deleted.

#### Mispicked Ion Merging

The algorithm identifies potential mispicked or misaligned [5–7] features using a narrow *m/z* (default ≤ 10 ppm) and RT (default ≤ 0.002 min) windows and with significant correlation (default $r \geq 0.80$). After graph construction, the feature with the highest intensity is retained and merged with other ions in family. With increased *m/z* tolerance, this filter may also help identify additional isotope-like duplicates. However, care should be taken to avoid incorrectly grouping true isomeric features. The filter implementation is conceptually similar to the *filter_mispicked_ions* function in the mpactr [5,6] package, and to *refineChromPeaks* in xcms [8]. This filter was not used in the present study and is recommended only when poor peak integration or feature misalignment is expected.

#### Saturated (Ringing) Ion Cleaning

The algorithm searches for saturated or 'ringing' artifacts associated with very high-abundance signals [5,6,9]. Features are sorted by increasing *m/z*, and candidate pairs are detected using a broad *m/z* window (default ≤ 0.95 Da), RT tolerance (default ≤ 0.05 min), intensity threshold for saturated signal (default > 1e5), intensity-ratio constraint (default 10 times higher for saturated signal), mass-difference direction (by default artifacts are strictly with higher mass, optionally could be bidirectional search), and high correlation (default $r \geq 0.80$). After graph construction, the most intense feature in each family is retained, while suspected saturated/ringing artifacts are removed. Previous work [9] suggests that saturated signal artifacts are mainly observed in TOF instruments and usually appear at higher *m/z* values than the saturated signal. However, this filter may also be used for refined isotope-like artifact search in bidirectional mode with adjusted parameters, such as a narrower *m/z* window of 0.05 Da and a lower anchor-intensity threshold of 1e4. Crucially, care should be taken to avoid incorrectly grouping true isomeric or co-eluting features. The filter implementation is conceptually related to *filter_mispicked_ions* function in mpactr package [6], and represents simplified version of the *rm.sat* function from the nontarget package [1] and the algorithm presented in the study [9]. We did not use it in the present study and suggest these filters only if expected poor integration or supersaturated signal.

### Orbi dataset

Detailed sample preparation and LC-MS analysis descriptions with raw data are available in [10].

#### Data Processing

Raw files were converted into mzML via MSConvert [11].

**MZmine**

The MZmine (4.8.0) [12] was run with UPLC-DDA workflow template; parameters were as follows:

| Module | Parameters used |
|---|---|
| Mass detection | Noise level MS1 = 5E2<br>Noise level MS2 = 1E2 |
| Chromatogram Builder | Scan filters:<br>level = 1<br>Minimum consecutive scans = 4<br>Minimum intensity for consecutive scans = 1E3<br>Minimum absolute height = 1E3<br>mz tolerance (scan-to-scan) = 0.005 Da or 20 ppm |
| Smoothing | Smoothing algorithm = Savitzky Golay<br>Retention time smoothing = 5 |
| Local minimum feature resolver | Dimension = Retention time<br>Chromatographic threshold = 90 %<br>Minimum search range RT/Mobility (absolute) 0.05<br>Minimum relative height = 0%<br>Minimum absolute height = 1E3<br>Min ratio of peak top/edge = 1.8<br>Peak duration range = 0.0 – 1.51<br>Minimum scans (data points) = 4 |
| 13C isotope filter | m/z tolerance = 0.0015 Da or 3 ppm<br>RT tolerance = 0.04 min<br>Maximum charge = 2 |
| Isotopic peaks finder | Chemical elements = H, C, N, O, S<br>m/z tolerance (feature-to-scan) = 0.0015 Da or 3 ppm<br>Maximum charge of isotope m/z = 1 |

| | |
|---|---|
| Join aligner | m/z tolerance (sample-to-sample) = 0.004 Da or 8 ppm<br>Weight for m/z = 3<br>Retention time tolerance 0.1 min<br>Weight for RT = 1 |
| Feature list rows filter | Sample-based filters:<br>Minimum aligned samples: 1 or 0.0 % |
| Feature finder | Intensity tolerance = 20 %<br>m/z tolerance = 0.005 Da or 20 ppm<br>Retention time tolerance = 0.1 min<br>Minimum scans (data points) = 2 |
| Duplicate peak filter | m/z tolerance = 0.0008 or 1.5 ppm<br>RT tolerance = 0.04 min |
| Correlation grouping (metaCorrelate) | RT tolerance = 0.06 min<br>Minimum feature height = 0<br>Intensity threshold for correlation = 5E2<br>Min samples in all: 1 or 0 %<br>Min samples in group 0 or 0 %<br>Min %-intensity overlap = 60 %<br>Exclude gap-filled features = True<br>Feature shape correlation:<br>Min data points = 5<br>Min data points on edge = 2<br>Measure = PEARSON<br>Min feature shape correlation = 85 %<br>Min total correlation = 50 %<br>Feature height correlation:<br>Minimum samples = 2<br>Measure = PEARSON<br>Min correlation = 70 % |
| Ion identity networking | m/z tolerance = 0.0015 Da or 3 ppm<br>Min height = 0<br>Default positive adducts list |
| Spectral / Molecular Networking | Algorithm = Modified cosine<br>m/z tolerance = 0.005 Da or 20 ppm<br>Merge (simple) |

Merging m/z tolerance = 0.005 Da or 20 ppm
Max precursor m/z delta = 500
Minimum matched signals = 4
Min cosine similarity = 0.7
Remove residual precursor m/z = 10
Crop to top N signals = 250
Signal threshold = 50
Intensity filter = 98 %

##### GNPS

A feature-based molecular network (FBMN) was created using the online workflow (SERVER:2026.08.27;WORKFLOW:2026.08.18) at GNPS 2 platform [13]. The data were filtered by enabling window and precursor filters. A parent mass tolerance of 0.02 Da and a MS/MS fragment ion tolerance of 0.02 Da were used to create consensus spectra. A network was created in which edges were filtered to have a cosine score of >0.6 and more than 4 matched peaks, while a maximum mass shift was set to 1999. Max component size was set to 100, and Top K was 10. The spectra in the network were then searched against GNPS spectral library. All matches kept between network spectra and library spectra were required to have a cosine similarity score of >0.6 and at least 4 matched peaks and returned top 1 match. Analog search was disabled.

In total, the spectral data and peak table, both before and after MetaboCensoR filtering, were subjected to FBMN and to Ion Identity Molecular Networking (IIMN) [14] by uploading "Supplementary Pairs" file generated by MZmine.

#### Data Availability

Links to the GNPS projects:

Raw Data FBMN: https://gnps2.org/status?task=43da3f82c33f43198f40cdc82837b87d

After App FBMN: https://gnps2.org/status?task=f686537881184bf08d45e499acf18c3d

Raw Data IIMN: https://gnps2.org/status?task=fde43183ef3940aa94985576117b7854

After App IIMN: https://gnps2.org/status?task=0c58cb476c80455f8b4c2f82472eb683

A full network was processed in Cytoscape (3.10.3) [15] and RCy3 [16]. Network nodes filling was defined by SIRIUS (6.3.3) [17] output, computed in default settings. A border color was defined by MetaboAnnotation [18] in comparison with target compounds library. Links to the Cytoscape projects:

Raw Data FBMN: RAW_FBMN.cys ; After App FBMN: APP_FBMN.cys ; Raw Data IIMN: RAW IIMN.cys ; After App IIMN: APP IIMN.cys

### Folate dataset

#### Human Cell Lines

The dataset was provided by Dr. Mariam Fokra, Dr. Nikita Sarvin, Dr. Tomer Shlomi (Technion, Israel). Reh (CRL-8286, ATCC, USA) human cell line that was isolated from tissue from an acute lymphocytic leukemia (ALL) patient was used as a wild type (WT) control cell line. Two types of mutations were introduced to the WT in FPGS gene by CRISPR/Cas9 system. One type is the FPGS gene point mutation (PM) at the position E115K [19] identified in clinical samples diagnosed with relapsed leukemia, which is characterized by reduced FPGS activity. Introduction of PM was controlled by Sanger sequencing and two single clones were eventually selected (SC 31 and 37, labeled PM-31 and PM-37). The other type is knockout (KO) of FPGS encoded gene. Procedure was controlled by gel-electrophoresis and two single clones were eventually selected (SC 6 and 22, labeled KO-6 and KO-22). Overall, 5 cell lines were examined: WT, PM-31/37, and KO-6/22. The cell line was cultured in Roswell Park Memorial Institute medium (RPMI-1640) supplemented with 10% dialyzed fetal bovine serum (FBS), 2 mM L-glutamine, 100 U/ml penicillin, and 100 mg/ml streptomycin, and with a physiological level of the folic acid [20] at 37 °C and 5% $CO_2$, passaging 3 times a week (no more than 15 passages) during all experiments. Cell line was tested for mycoplasma using EZ-PCR mycoplasma detection kit (Biological Industries USA, Inc., USA). Cell counting was performed by Beckman Coulter Z2 Cell and Particle Counter (Beckman Coulter, USA) for the cell size from 7 to 27.55 um and according to the manufacturer's instructions.

#### Sample Preparation

First, cells were collected into a microtube, counted and pelleted via centrifugation at 500 g for 5 min at 4°C. After washing two times with ice-cold PBS, cell pellets were resuspended in appropriate volume of MeOH–CH3CN–$H_2O$ (5:3:2 vol/vol/vol) extraction mixture (sample-to solvent ratio 25 mln cells/ml), homogenized at vortex for 1 min, spun down, and froze in liquid N2 for 15 min. Frozen extracts were kept at -80°C before analysis and for at least 2 hours. Prior analysis, extracts were defrosted for 5 min at room temperature, vortexed again and centrifuged twice at 14 000 rpm (0°C, 20 min) and then analyzed using LC-MS. Collected samples were analyzed together with pooled QC samples, which were injected after each 5 samples. All LC-MS runs were acquired during a single batch and in blocked design to prevent systematic bias.

#### LC-MS Analysis

Metabolomics extracts were analyzed with Thermo Scientific UltiMate 3000 HPLC system coupled with Thermo Scientific Q Exactive Orbitrap. Chromatographic separation was achieved on a SeQuant ZIC-pHILIC column (2.1 × 150 mm, 5 µm, EMD Millipore, USA) with guard column. Flow rate was set to 0.2 $mL \times min^{-1}$, column compartment was set to 30 °C, and autosampler tray was maintained at 4 °C. Mobile phase A consisted of 20 mM ammonium carbonate and 0.01% (v/v) ammonium hydroxide. Mobile Phase B was 100% acetonitrile. The mobile phase linear gradient (%B) was as follows: 0 min 80%, 12.5 min 30 %, 15.0 min 30%, 15.2 min 80%, 23.0 min 80%. Injection volume was 5 uL. Mass detection has been performed with an ESI ionization source working in negative ionization mode. Ionization source parameters were as follows: sheath gas 25 units, auxiliary gas 3 units, spray voltage 3.3 kV, capillary temperature 325 °C, S-lens RF level 65, auxiliary gas temperature 200 °C. Accumulation time was set to 150 ms, resolution - 70000; automatic gain control - $3.00E^{+06}$. Metabolites were analyzed in the range 72–1080 *m/z*. All LC-MS runs were acquired during a single batch and in blocked design to prevent systematic bias.

#### Data Processing

Raw files were converted into mzXML via MSConvert [11].

**El-Maven**

Because FPGS enzyme is involved in folate metabolism, we expected pathway-level impacts related to folate-dependent processes, nucleotide, and energy pathways, including: Energy metabolism; Pentose Phosphate Pathway; Purine; Pyrimidine; SAM, Gly-Ser. For manual examination of affected pathways peak integration was performed by EI-Maven [21] (0.12.0) software by constructing EIC based on molecular formula for metabolites involved in selected pathways according to the HumanCyc [22] database and with assumption of formation $[M\text{-}H]^-$ adduct. Retention time for 48 metabolites was confirmed by injection of standard compound.

**xcms**

The xcms (3.17.3) [8] parameters optimized in IPO (1.20.0) [23] were as follows:

| XCMS functions: | Parameter |
|---|---|
| findChromPeaks | CentWaveParam:<br>ppm = 14.9<br>mzdiff = -0.00265<br>snthresh = 10 |

| | |
|---|---|
| | noise = 1135<br>prefilter = 3, 146.46<br>mzCenterFun = wMean<br>integrate = 1<br>fitgauss = FALSE |
| adjustRtime | obiwarpParam:<br>binSize = 1<br>response = 18<br>distFun = cor_opt<br>gapInit = 0.5<br>gapExtend = 3<br>factorDiag = 2<br>factorGap = 1<br>localAlignment = FALSE |
| groupChromPeaks | PeakDensityParam:<br>bw = 22<br>minFraction = 0.9<br>binSize = 0.001<br>maxFeatures = 50 |
| fillChromPeaks | default settings |

#### Statistical Analysis

Prior univariate analysis, zero values were replaced by the randomly generated values from the pre-defined distribution (μ = noise level, SD = 0.3×noise level). T-test p-values were adjusted for multiple testing using the Benjamini–Hochberg FDR procedure. All computations were performed in the R environment (version 4.1.2) [24].

#### Data Availability

The converted raw data is available in the MassIVE repository under the accession number: MSV000100951 (https://doi.org/10.25345/C51J97N5K).

### Inter dataset

#### Bacterial Strains

Study involved *Paenibacillus dendritiformis* (*Pd*), *Bacillus subtilis* NCIB 3610 (*Bs*, WT) and the NRPS-mutated *dsrf*, *dpps*, and *dd* that are Surfactin and/or Plipastatin Synthetase deficient.

A suspension of bacteria (10 μL) was inoculated from overnight cultures in LB liquid medium for microbial interactions on the nutrient plate with 0.5% bacto peptone and 1% agar. Plates were incubated at 30 °C for 24 h before analysis.

*Pd* was co-inoculated with itself, other bacteria, or 0.01 mg of commercial Surfactin at 1 cm distance to study interactions [25].

#### Sample Preparation

The agar in each plate was cut in half following dividing by several regions, and each region was further cut into small pieces using a sterile surgical blade and then transferred into the tube. The wet weight of each sample was used to normalize the solvent-sample ratio, with 100 μL of cold MeOH (-20°C) added per 100 mg of sample. Sample was vortexed for 1 minute, spun down, and left on the bench overnight. Then it was sonicated in an ultrasonic bath at 40 kHz for 90 minutes, with the water changed every 45 minutes. After mixing, the sample was precipitated by centrifugation at 20 000 g for 20 min at 4°C and 200 μL of supernatant was taken to a new tube and mixed with equal water volume. After mixing, resulted extract was precipitated by centrifugation at 20 000 g for 20 min at 4°C and supernatant was transferred into the amber glass vial for subsequent analysis.

#### LC-MS Analysis

Samples were analyzed with Thermo Scientific Vanquish HPLC system coupled with Bruker TIMS-TOF Pro 2 mass spectrometer. Chromatographic separation was achieved on a Kinetex C18 column (2.1 × 50 mm, 1.7 μm, Phenomenex, USA) with guard column. Flow rate was set to 0.3 $mL \times min^{-1}$, column compartment was set to 35 °C, and autosampler tray was maintained at 4 °C. Initial mobile phase A consisted of water with 0.1% (v/v) formic acid, mobile phase B was acetonitrile with 0.1% (v/v) formic acid. The mobile phase linear gradient (%B) was as follows: 0.0-1.0 min 5 %, 7.0 min 95 %, 9.0-10.5 min 95%, 11.0 min 5 %, 15.0 min 5 %. The injection volume was 5 uL. Mass detection has been performed with an ESI ionization source working in positive mode. Ionization source parameters were as follows: 500

V end plate, 4500 V capillary voltage, 2.2 Bar nebulizer, dry gas 10 l/min, dry temperature 220°C. Detection was performed in SCAN and DDA (Auto MS/MS) modes simultaneously. Metabolites were recorded in the range 90-2500 *m/z*. DDA parameters were as follows: MS spectra rate 1 Hz, MS/MS spectra rate 12 Hz, MS/MS spectra rate limits 12-20 Hz, rate control: Dynamic, total cycle time 1.83 s, target intensity 10 000, number of precursors: 10, active exclusion after 3 spectra and release after 1 min, normalized threshold: 200 counts / 1000 scans. MS/MS acquisition used charge- and precursor *m/z*-dependent collision energies. For singly charged ions, CE increased from 20 to 70 eV across the 50–2000 *m/z* range, while doubly and triply charged ions were fragmented using 35–50 eV, with isolation widths of 2–15 *m/z*. All LC-MS runs were acquired during a single batch and in blocked design to prevent systematic bias.

#### Data Processing

Raw files were converted into mzXML via MSConvert [11].

**MS-DIAL**

The MS-DIAL [26] (5.5.251021) parameters were as follows:

| Step | Parameters |
|---|---|
| Data Collection | MS1 tolerance = 0.001 Da |
| | MS2 tolerance = 0.005 Da |
| | Retention time begin = 0 min |
| | Retention time range end = 100 min |
| | Maximum charge number = 2 |
| Peak Detection | Minimum peak height = 10,000 |
| | Mass slice width = 0.05 Da |
| | Smoothing method = Linear Weighted Moving Average |
| | Smoothing level = 3 |
| | Minimum peak width = 5 |
| Spectrum deconvolution | - |
| Adduct Ion | Adducts: all available positively charged adducts |
| Alignment parameters | Reference file = file ending 3862 |
| | Retention time tolerance = 0.2 min |
| | MS1 tolerance = 0.003 Da |
| | Retention time factor = 0.5 |
| | MS1 factor = 1 |
| | Peak count filter = 0% |

N% detected in at least one group = 0%
Gap filling by compulsion = True

#### Statistical Analysis

Prior univariate analysis, zero values were replaced by the randomly generated values from the pre-defined distribution ($\mu$ = noise level, SD = 0.3×noise level). T-test p-values were adjusted for multiple testing using the Benjamini–Hochberg FDR procedure. All computations were performed in the R environment (version 4.1.2) [24].

To evaluate the impact of data filtering on statistical power (Pd+Pd vs. Pd+Bs), effect sizes were calculated as Hedges' g (95% CI) using the effectsize R package [27]. Absolute power was then estimated using the pwrFDR package [28] for a balanced design (n=4 per group, alpha=0.05) with Benjamini-Hochberg FDR control. To account for dimensionality reduction, the total number of tests in the power estimation was set to 6,027 for the raw dataset and 509 for the MetaboCensoR-filtered dataset.

#### Data Availability

The converted raw data is available in the MassIVE repository under the accession number MSV000100949 (https://doi.org/10.25345/C5930P825).

#### Supplementary Figs. 1-2. Mass spectrum annotation of unreported degradation product

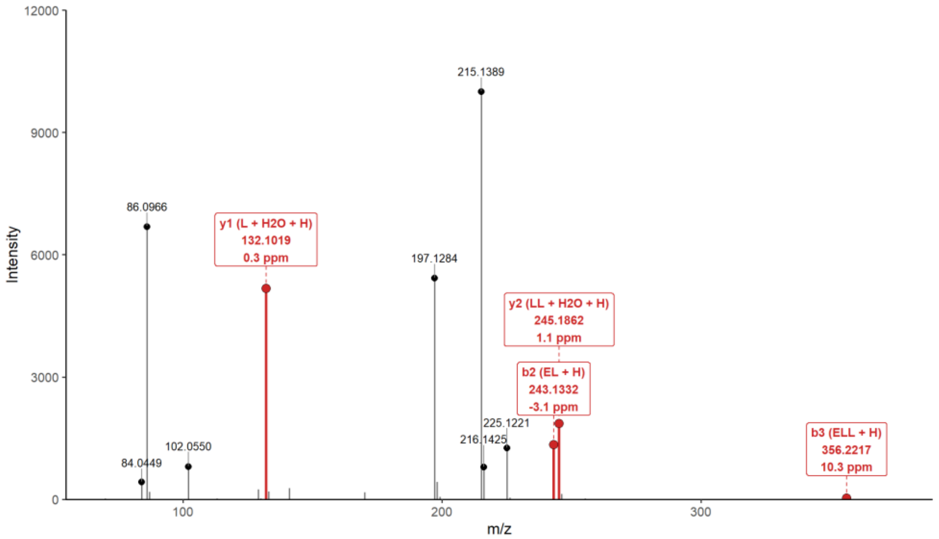

Figure S1. Putative MS/MS annotation of the previously unreported peptidic degradation product (C17H31N3O6; Glu–Leu–Leu; parent ion *m/z*: 374.2294; $[M+H]^+$; RT 4.87 min). Mass spectrum is available in .ms and .mgf formats from MassIVE repository under the accession number MSV000100949.

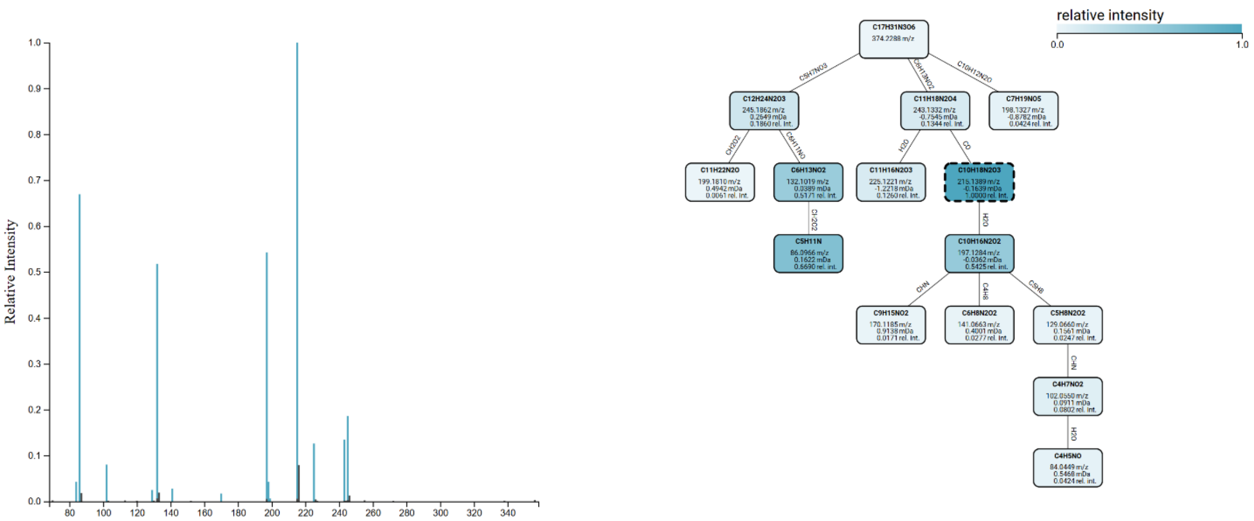

Figure S2. Mass spectrum annotation in SIRIUS [17] of fragmentation spectra for previously unreported peptidic degradation product (C17H31N3O6; Glu–Leu–Leu; parent ion *m/z*: 374.2294; $[M+H]^+$; RT 4.87 min). Mass spectrum is available in .ms and .mgf formats from MassIVE repository under the accession number MSV000100949.

### Benchmarking Parameters

#### khipu / MetaboCensoR

An mzMine-generated peak table from the orbi dataset, after blank filtering, was used as input for both tools. mzMine settings were identical to those described for the orbi dataset.

khipu [29,30] was run using its online implementation (https://metabolomics.cloud/khipu/process) with mass and RT tolerances of 10 ppm and 0.3 s, respectively.

For MetaboCensoR, MS filtering was performed using the default settings for isotope and adduct collapsing.

#### mpactR / MetaboCensoR

The mzMine-generated peak table from the orbi dataset was used as input for both tools.

mpactR (version 0.2.1) [5,6] was applied sequentially using mispicked-ion filtering (ringwin = 0.5, isowin = 0.01, trwin = 0.005, max_iso_shift = 3, merging = "sum"), blank filtering (threshold = 0.1), CV filtering (threshold = 0.3), and in-source fragment filtering (cluster = 0.95).

Corresponding filtering steps were applied in MetaboCensoR: blank filtering (threshold = 0.1), isotope and adduct collapsing, in-source fragment filtering with MS2 matching (0.01 rt threshold) and with all other parameters kept at their default values, and RSD filtering (threshold = 30%).

#### MS-CleanR / MetaboCensoR

Because MS-CleanR is compatible only with MS-DIAL 4, an MS-DIAL-generated peak table from the orbi dataset was used for this comparison. MS-DIAL (version 4.12) was configured according to the settings described in the MS-CleanR manuscript [31]: MS1 and MS2 tolerances of 0.01 and 0.05 Da, respectively, centroid data, a minimum peak height of 10,000, and alignment to sample WS03B using RT and mass tolerances of 0.1 min and 0.015 Da, respectively.

MS-CleanR (version 1.0) was applied sequentially using blank filtering (ratio = 0.8), incorrect-mass filtering, ghost-peak filtering, RSD filtering (30%), and RMD filtering (50–3000). Within each ion family, two features were retained based on intensity and network connectivity.

Corresponding steps were applied in MetaboCensoR, including blank filtering (threshold = 0.1), isotope and adduct collapsing, in-source fragment filtering with MS2 matching (0.01 rt threshold) and with all other parameters kept at their default values; RSD filtering (threshold = 30%), and RMD filtering (50–3000). Because MS-CleanR considers only features with acquired MS/MS spectra, the resulting MetaboCensoR-filtered table was additionally restricted to features with MS/MS data before comparison.

##### Binner / MetaboCensoR

An mzMine-generated and blank-filtered peak table from the orbi dataset was used as input for both tools.

Binner (version 1.0.0) [32] was run using RT and mass tolerances of 0.01 min and 0.005 Da for deisotoping and 0.005 min and 0.005 Da for annotation. Feature cleaning additionally included CV filtering (threshold = 4).

Corresponding filtering steps were applied in MetaboCensoR: isotope and adduct collapsing, in-source fragment filtering with MS2 matching (0.01 rt threshold) with all other parameters kept at their default values, and RSD filtering (threshold = 400%, equal to CV = 4).

[14] R. Schmid, D. Petras, L.F. Nothias, M. Wang, A.T. Aron, A. Jagels, H. Tsugawa, J. Rainer, M. Garcia-Aloy, K. Dührkop, A. Korf, T. Pluskal, Z. Kameník, A.K. Jarmusch, A.M. Caraballo-Rodríguez, K.C. Weldon, M. Nothias-Esposito, A.A. Aksenov, A. Bauermeister, A. Albarracin Orio, C.O. Grundmann, F. Vargas, I. Koester, J.M. Gauglitz, E.C. Gentry, Y. Hövelmann, S.A. Kalinina, M.A. Pendergraft, M.

Panitchpakdi, R. Tehan, A. Le Gouellec, G. Aleti, H. Mannochio Russo, B. Arndt, F. Hübner, H. Hayen, H. Zhi, M. Raffatellu, K.A. Prather, L.I. Aluwihare, S. Böcker, K.L. McPhail, H.U. Humpf, U. Karst, P.C. Dorrestein, Ion identity molecular networking for mass spectrometry-based metabolomics in the GNPS environment, Nat. Commun. 12 (2021) 3832. https://doi.org/10.1038/s41467-021-23953-9.

[22] M. Trupp, T. Altman, C.A. Fulcher, R. Caspi, M. Krummenacker, S. Paley, P.D. Karp,

Beyond the genome (BTG) is a (PGDB) pathway genome database: HumanCyc, Genome Biol. 11 (2010) O12. https://doi.org/10.1186/gb-2010-11-s1-o12.
[23] G. Libiseller, M. Dvorzak, U. Kleb, E. Gander, T. Eisenberg, F. Madeo, S. Neumann, G. Trausinger, F. Sinner, T. Pieber, C. Magnes, IPO: A tool for automated optimization of XCMS parameters, BMC Bioinformatics 16 (2015) 118. https://doi.org/10.1186/s12859-015-0562-8.
[24] R Core Team, A Language and Environment for Statistical Computing., R Found. Stat. Comput. (2021). https://www.r-project.org/.
[25] T. Luzzatto-Knaan, A. V. Melnik, P.C. Dorrestein, Mass Spectrometry Uncovers the Role of Surfactin as an Interspecies Recruitment Factor, ACS Chem. Biol. 14 (2019) 459–467. https://doi.org/10.1021/acschembio.8b01120.
[26] H. Takeda, Y. Matsuzawa, M. Takeuchi, M. Takahashi, K. Nishida, T. Harayama, Y. Todoroki, K. Shimizu, N. Sakamoto, T. Oka, M. Maekawa, M.H. Chung, Y. Kurizaki, S. Kiuchi, K. Tokiyoshi, B. Buyantogtokh, M. Kurata, A. Kvasnička, U. Takeda, H. Uchino, M. Hasegawa, J. Miyamoto, K. Tanabe, S. Takeda, T. Mori, R. Kumakubo, T. Tanaka, T. Yoshino, M. Okamoto, H. Takahashi, M. Arita, H. Tsugawa, MS-DIAL 5 multimodal mass spectrometry data mining unveils lipidome complexities, Nat. Commun. 15 (2024) 9903. https://doi.org/10.1038/s41467-024-54137-w.
[27] M. Ben-Shachar, D. Lüdecke, D. Makowski, effectsize: Estimation of Effect Size Indices and Standardized Parameters, J. Open Source Softw. 5 (2020) 2815. https://doi.org/10.21105/joss.02815.
[28] G. Izmirlian, Strong consistency and asymptotic normality for quantities related to the Benjamini–Hochberg false discovery rate procedure, Stat. Probab. Lett. 160 (2020) 108713. https://doi.org/10.1016/j.spl.2020.108713.
[29] S. Li, S. Zheng, Generalized Tree Structure to Annotate Untargeted Metabolomics and Stable Isotope Tracing Data, Anal. Chem. 95 (2023) 6212–6217. https://doi.org/10.1021/acs.analchem.2c05810.
[30] J.M. Mitchell, Y. Chi, S. Zheng, M. Thapa, E. Wang, S. Li, Annotation of Metabolites in Stable Isotope Tracing Untargeted Metabolomics via Khipu-web, J. Am. Soc. Mass Spectrom. 35 (2024) 2824–2835. https://doi.org/10.1021/jasms.4c00175.
[31] O. Fraisier-Vannier, J. Chervin, G. Cabanac, V. Puech, S. Fournier, V. Durand, A. Amiel, O. André, O.A. Benamar, B. Dumas, H. Tsugawa, G. Marti, MS-CleanR: A Feature-Filtering Workflow for Untargeted LC-MS Based Metabolomics, Anal. Chem. 92 (2020) 9971–9981. https://doi.org/10.1021/acs.analchem.0c01594.

[32] M. Kachman, H. Habra, W. Duren, J. Wigginton, P. Sajjakulnukit, G. Michailidis, C. Burant, A. Karnovsky, Deep annotation of untargeted LC-MS metabolomics data with Binner, Bioinformatics 36 (2020) 1801–1806. https://doi.org/10.1093/bioinformatics/btz798.
